# miRNA synergism impaired metabolic programming of oral cancer cells by selectively enhancing target gene specificity

**DOI:** 10.1101/2024.08.01.606174

**Authors:** Sayani Mazumder, Urbee Banerjee, Simran Sinsinwar, Saubhik Mandal, Debashis Prusty, Prabhakar Babele, Bhabatosh Das, Soumen Manna, Jay Gopal Ray, Raghunath Chatterjee

## Abstract

Despite the emergence of miRNAs as promising therapeutic tools in cancer management, most clinical trials have not been successful due to their cytotoxic effects. Here, we have investigated the factors regulating the effect of miRNA-mimic pairs in enhancing target gene specificity while reducing cellular toxicity. Synergistic reductions by the miRNA-mimic pairs were observed for the 3’-UTRs of common target genes with miRNA-responsive elements (MREs) preferentially located at a distance of 200-800bp. Deletion of either of the miRNA seed sequences resulted in a loss of synergism. Furthermore, we performed small RNA-sequencing to identify significantly downregulated miRNAs in Oral cancer. Consequently, we transfected let-7c-5p and miR-125b-5p miRNA mimics either alone or in combination at half-dose concentrations and determined the expression levels of their common and unique target genes in oral cancer cells. Significant reductions in target gene expression were observed for genes containing MREs for both miRNA mimics within the preferential distance. Proteomic data revealed that the let-7c-5p and miR-125b-5p miRNA-mimic pairs synergistically reduced the expression of their common target genes, hexokinase 2 (HK2) and branched-chain amino acid transaminase 1 (BCAT1), in cancer cells. However, unique target genes did not exhibit any significant alterations in their protein levels. The effect of HK2 and BCAT1 downregulation was also reflected in the metabolomic profiling of cancer cells, specifically affecting glycolysis and the BCAA degradation pathways. As a result of the metabolic impairment caused by the synergistic effect of miRNA mimic pairs, oral cancer cells showed a significant reduction in proliferation, migration, and spheroid formation compared to cells treated with either single miRNA. The synergistic effect of these miRNA mimics was lost in normal keratinocytes, possibly due to low expression levels of the target oncogenes, suggesting a cancer cell-specific effect of this mechanism.

## 1. INTRODUCTION

MicroRNAs (miRNAs) are a class of small noncoding RNAs that regulate gene expression by inducing translational repression or mRNA decay [1, 2]. MiRNA deregulation in many cancers occurs through a variety of mechanisms, including mutation, deletion, amplification, and epigenetic silencing [3]. MiRNAs possess therapeutic potential either as standalone treatments or in combination with other therapeutic approaches [4, 5]. A double-stranded miR-34 mimic, enclosed within a liposome-formulated nanoparticle (MRX34), was the first miRNA used in therapeutics [4]. Elevated levels of miRNA-34a in cancer have been shown to diminish the proliferative capacity of cells [6]. However, the first trials of miRNA therapy using MRX34 were unsuccessful and terminated due to severe immune-mediated adverse effects [4, 7]. Another drug, TargomiR, a synthetic double-stranded miR-16 mimic, had a moderate effect, possibly due to the low dose of the mimic and minor adverse effects [7, 8]. Clinical trials with inhibitors of miR-155, an abundantly expressed miRNA in lymphomas, demonstrated the elimination of cutaneous lesions [9, 10]. Similarly, *in vivo* delivery of anti-miR-10b significantly reduced metastasis in breast carcinoma [11]. The combination of miRNA with other antitumor modalities has been documented to target a wider spectrum of tumors, enhance therapeutic effectiveness, and overcome drug resistance [5]. The coadministration of a miR-21 inhibitor, coupled with the chemotherapeutic drug doxorubicin, demonstrated notable anticancer efficacy[12]. Similarly, a combination of miR-29b and bortezomib, used to treat multiple myeloma, revealed that miR-29b could enhance cancer cells’ sensitivity to bortezomib [13].

Despite these advancements, miRNA-mediated cancer therapy has not been successfully employed as an effective treatment strategy, primarily because of the cytotoxic effects of higher doses of miRNA mimics/anti-miRs required for effective treatment. To date, only two clinical trials of microRNAs for the treatment of cancers have been approved, and both have shown partial responses to the treatment with evidence of miRNA induced toxicity [8, 14]. To overcome these problems, a reasonable approach could be to use lower dose combinations of miRNAs that synergistically regulate the expression of a mutual target [15]. Recently, the synergistic effect of miRNAs in cancer therapeutics has gained interest [16, 17]. Young *et al.* (2013) studied the potential importance of the secondary structures of pre-miRNAs in both the cooperativity and specificity of target genes [16]. Schmitz *et al.* (2014) also reported that miRNA cooperativity is a frequent mechanism for enhanced target repression by miRNA pairs [17]. Noguchi *et al.* (2013) studied the antitumor effects of combination treatment with miR-143 and miR-145 on bladder cancer cell lines and reported that the ectopic expression of both miRNAs led to significant synergistic growth inhibition [18]. Similarly, the synergistic effect of miR-34a and miR-15a/16 induces cell cycle arrest in an Rb-dependent manner in non-small cell lung carcinoma [19]. MiR-124 and miR-203 have also been reported to synergistically inactivate the EMT pathway in clear cell renal cell carcinoma (ccRCC) by targeting ZEB2 [20].

MiRNA-mediated synergism might depend on several underlying factors, such as the distance between the MREs, Gibb’s free energy, and the selection of appropriate miRNAs and their target genes. Previous reports have suggested that a distance of 13-35nt between MREs in the 3’-UTR sequence of their target genes is associated with co-operativity [21]. In contrast, another study showed an antagonistic effect with the co-binding of the miRNA-pairs on the 3’-UTR of the same target gene, while a synergistic effect was reported with miRNA-pair binding in the 3’-UTRs of different genes or in the 5’-UTR of the same target gene [22]. Although these studies have elucidated the importance of the distance between MREs for miRNA co-operativity, they have used synthetic constructs with a maximum distance up to 70 nt between MREs [21]. The effect of miRNA coexpression with varying and wider range of distances between the MREs with the UTRs of real target genes is still an open area of research and is largely unexplored. Moreover, no systematic studies have investigated the effect of miRNA-mediated synergism on off-target effects and cellular toxicity.

In the present study, we identified downregulated tumor-suppressor miRNAs and their inversely correlated target oncogenes in oral cancer. Instead of studying miRNA-mediated synergism using synthetic 3’-UTR constructs with specific ranges of distances between MREs, we utilized the native 3’-UTRs of the overexpressed oncogenes targeted by a pair of downregulated tumor suppressor miRNAs to examine the effect of the distance between MREs on miRNA-mediated synergism. Finally, we investigated the synergistic effects of miRNAs on off-target genes and cellular toxicity in oral cancer by profiling proteomic, metabolomic, and phenotypic assays.

## 2. MATERIAL & METHODS

### 2.1 Patient Selection and Sample Collection

Patients clinically diagnosed with oral squamous cell carcinoma (OSCC) were recruited from the eastern region of India. Only patients who provided informed written consent were included in the study. Tissue was excised from the lesion site and an adjacent normal area for each patient, preserved in RNAlater (Invitrogen), and stored at -80°C until further processing. Four patients were included in the discovery cohort, and 33 patients were included in the validation cohort following histopathological analysis and confirmation of well-differentiated carcinoma. All the tissue samples were collected from Dr. R. Ahmed Dental College and Hospital. The study was approved by the Institutional Ethics Committee for Human Research at the Indian Statistical Institute, Kolkata. The mean age of patients in the discovery cohort was 47.75 years, while the mean age in the validation cohort was 54.4 years **(Tables S1 and S2**).

### 2.2 Cell-lines and Plasmids

Luciferase assays and lentiviral particle generation were performed in HEK-293T cells (ATCC). DMEM media supplemented with GlutaMAX (Gibco) was used to culture the HEK-293T and HaCaT cell lines (AddexBio). The media was supplemented with 10% fetal bovine serum (FBS) (Gibco) and 1% penicillin-streptomycin (Pen-Strep) (Gibco). The ORL-48 oral cancer cell line was a generous gift from Prof. Sok Ching Cheong, Department of Oro-Maxillofacial Surgery and Medical Sciences, Faculty of Dentistry, University of Malaya, Kuala Lumpur, Malaysia. DMEM/F12 (1:1) media (Gibco) supplemented with 10% FBS, 1% Pen-Strep, and 0.5 μg/mL hydrocortisone (Gibco) was used to culture ORL-48 cells. These cell lines were subsequently used for the functional characterization of miRNAs in oral cancer. The cells were incubated at 37°C in a humidified atmosphere with 5% CO2.

The pRNA-U6 vector (GenScript) was used for the transient overexpression of miRNAs. The pLKO.1-TRC cloning vector (Addgene), psPAX2 (Addgene), and pMD2.G (Addgene) plasmids were used for constitutive overexpression of miRNAs using lentiviral transduction for generation of the miRNA overexpressing stable cells.

### 2.3 RNA Extraction

Total RNA was extracted from OSCC and adjacent normal tissue samples using RNeasy Mini Kit (Qiagen). Briefly, 10 µL of β-mercaptoethanol was freshly mixed with RLT Plus buffer (1 mL) before use. Five to ten milligrams of biopsy specimens collected in RNAlater were snap-frozen in liquid nitrogen, ground to a fine powder using a mortar and pestle, and RNA isolation was performed following the manufacturer’s protocol (http://www.qiagen.com/HB-0435-007). The isolated RNA was kept on ice during downstream processing and stored at -80°C. A Nanodrop 2000 Spectrophotometer (Thermo Scientific) and Bioanalyzer (Agilent) were used to check the quantity and quality of the eluted RNA.

### 2.4 Illumina Small RNA Sequencing

Genome-wide small RNA-sequencing of 4 OSCC tissues and 4 adjacent normal tissues was performed. One microgram of total RNA was used as an input for library preparation using Illumina TruSeq Small RNA Library Preparation Kit following the manufacturer’s protocol (TruSeq Small RNA Library Prep Reference Guide), followed by sequencing on an Illumina HiSeq 2500 instrument. The TruSeq SBS Kit v3 was used for single-end sequencing. Following RNA-sequencing, the qualities of the sequenced reads were assessed using FastQC [23], followed by 3’ and 5’ adapter trimming using Cutadapt [24]. Reads with a minimum length of 16 bases were aligned to the reference human genome (GRCh37/hg19) (https://genome.ucsc.edu/) using NovoAlign software in miRNA mode (https://www.novocraft.com/). The raw counts for each annotated miRNA were obtained using HTSeq-Count [25] followed by differential expression analysis using EdgeR[26]. A False Discover Rate (FDR) adjusted p-value <0.05 was used as the cutoff for identification of significantly differentially expressed miRNAs between OSCC tissues and adjacent normal tissues. For OSCC, miRNAs with a Log(FC) ≥1, Log(CPM) ≥2 and FDR ≤0.05 were considered as upregulated while miRNAs with a Log(FC) ≤1, Log(CPM) ≥2 and FDR ≤0.05 were considered as downregulated in OSCC (**Table S3**).

### 2.5 Comparative Study: Small RNA Sequencing vs. TCGA-HNSC Small RNA Sequencing

The small RNA-sequencing data were compared with the small RNA-sequencing dataset of HNSCC from TCGA. The publicly available “The Cancer Genome Atlas (TCGA)” small RNA-sequencing data of HNSCC patients were downloaded from TCGA-GDC Data Portal (NIH) (https://portal.gdc.cancer.gov). TCGA-HNSC small RNA-sequencing data included 30 normal samples, 20 Stage-I patients, 54 Stage-II patients, 61 Stage-III patients and 164 Stage-IV patients. The manifest files and the clinical data files for each patient were downloaded using the GDC Data Transfer tool (https://github.com/NCI-GDC/gdc-client). The raw read counts were extracted and normalized, and the HTseq count[25] was used to count the number of reads mapped. EdgeR[26] was used for analysis of the statistically significant differentially regulated miRNAs. In both datasets, miRNAs with a |Log (FC)|≥1, Log (CPM)≥2 and FDR≤0.05 were considered differentially regulated in disease. The miRNAs that were found to be significantly deregulated in any one or more stages in TCGA dataset and our small RNA-sequencing dataset were considered common differentially regulated miRNAs across different populations.

### 2.6 Survival and KEGG Pathway Analysis

To determine the clinical relevance of the differentially expressed miRNAs, survival analysis was performed. The follow-up data and expression values for the differentially expressed miRNAs for each of the TCGA patients, available from TCGA website, were collected and used for survival analysis. Survival analysis of TCGA patients via the miRNA expression profile was performed with the OncoLnc online tool [27]. For the survival analysis, we considered the top and bottom 25^th^ percentiles of expressed miRNAs as the high and low groups, respectively. A result was considered to be significant if the Mantel-Cox log-rank p value was less than or equal to 0.05.

The union of genes targeted by the upregulated and downregulated miRNAs was identified from the microT-CDS database, and a KEGG pathway analysis was performed using miRPathv.3 (Diana tools) [28] to identify the top enriched pathways targeted by them. A p-value≤0.01 and a microT threshold of 0.8 were used as cutoff values. The union of genes targeted by miR-125b-5p, miR-204-5p and Let-7c-5p was also extracted from the microT-CDS database, and the top enriched pathways targeted by the union of genes targeted by these miRNAs were identified using the KEGG pathway.

### 2.7 Reverse Transcription and Quantitative Real-time PCR

One microgram of total RNA was used as a template for complementary DNA (cDNA) synthesis using the miScript II RT Kit according to the manufacturer’s protocol (http://www.qiagen.com/HB-0235). Ten nanograms of cDNA was used as a template for quantitative real-time PCR (qPCR) using QuantiTect® SYBR® Green PCR master mix (Qiagen). Customized 96-well array plates in which the miRNA specific forward primers were embedded in the wells were used for the expression study. A total of 1umole of universal reverse primer, 5X SYBR Green and 10ng of cDNA were used in a total reaction volume of 25uL. The PCR mixture was subjected to the following conditions: 95°C for 10 minutes, followed by 40 cycles at 95°C for 15 seconds, 55°C for 30 seconds and 72°C for 30 seconds. The nonspecific PCR products were checked by running the dissociation protocol. RNA-U6 was used as an endogenous control while miRTC and PPC were used as reverse transcription control and PCR reaction controls, respectively. A no-template control (NTC) was included on each experimental plate. All qPCR reactions were carried out on an ABI PRISM 7900 HD Detection System (Applied Biosystems).

### 2.8 Overexpression of miRNAs

The pre-miRNA sequences were cloned into the pLKO.1 vector (Adgene) by double digestion with Age-I (NEB) and EcoR-I (NEB) followed by transformation into *E.coli* Stbl3 cells. One microgram of the transformed pLKO.1 plasmid was cotransfected in HEK-293T cells with 750ng psPAX2 and 500ng pMD2.G vectors (Adgene) using Lipofectamine 2000 (Thermo Fisher) following manufacturer’s protocol (Protocol Pub. No. MAN0007824 Rev.1.0). The virus particles were harvested after 24 hours of transfection and 100µL of the harvested viral titer was used for infection to ORL-48 cell lines followed by puromycin selection. The stable cells constitutively overexpressing the miRNAs or scrambled DNA (ORL48_Let7c, ORL48_miR125b, ORL48_miR204, and ORL48_scrm) were verified by qPCR.

For transient overexpression of miRNA mimics, cells were seeded to reach 70-80% confluence after 24hour, and transient transfection of miRNA mimics (20pmol) was conducted using Lipofectamine RNAiMAX according to the manufacturer’s protocol (Protocol Pub. No. MAN0007825 Rev.1.0). For transient overexpression of miRNA mimics, cells were seeded to reach 70-80% confluence after 24hour and transient transfection of miRNA mimics (20pmol) was conducted using Lipofectamine RNAiMAX according to the manufacturer’s protocol.

### 2.9 Phenotypic Assays

#### 2.9.1 WST1 Cell Proliferation Assay

Cells were seeded in a 96-well plate (0.5×10^4^ - 2×10^4^ cells) and allowed to reach 70% confluency within 24hour. The following day, the media was discarded, 100μL complete media containing 10μL of WST1 reagent (Roche) was added to each well, and the cells were incubated at 37°C for 30-45min. The absorbance was measured at 450nm using an iMark Microplate Reader (Bio-Rad). The OD values of the stable cells or the transfected cells were normalized to the OD values of the control cell-line to calculate the proliferation rate.

#### 2.9.2 Wound-healing Cell Migration Assay

Cells were seeded in 35mm cell-culture dishes to reach 100% confluence. A scratch was made along the diameter of the dish, and the cells were washed two times with 1X dPBS, followed by the addition of 2mL of complete media. Wound closure was measured at regular intervals (0-hour, 3-hour, 6-hour, 9-hour) using an inverted microscope (Leica DMi1). The wound closure distances at different time points were normalized to the distance at 0hour to calculate the migration rate.

#### 2.9.3 FITC-Annexin V/PI Apoptosis Assay

The cells were seeded in 60 mm cell culture dishes and allowed to reach 70-80% confluence within 24hour. Apoptosis assay was performed using a Dead Cell Apoptosis Kit with Annexin V Alexa FluorTM 488 & propidium iodide (PI) (Invitrogen). The cells were counted and resuspended in 1mL of complete media at a dilution of 1×105 cells/mL. The samples were washed with 1mL of dPBS and resuspended in 100 μL of 1X binding buffer solution. Two microliters of Annexin-FITC conjugate and 1μL of PI were added to the cell suspension, which was subsequently vortexed and incubated for 15 min at RT in the dark. Finally, 400μl of 1X binding buffer was added and analyzed using a flow cytometer (FACSVerse, BD Biosciences, USA). Using FlowJO software v.9.0 (BD Biosciences, USA), the cell populations were selected by gating of t h e P1 and P2 populations and then analyzed to determine the fluorescence intensity in both the FITC and PI channels.

#### 2.9.4 Spheroid Formation Assay

Cells were seeded in a 24-well cell-culture dish and allowed to reach 70-80% confluence within 24hour. Transient transfection of miRNA mimics (20pmol) was conducted using Lipofectamine RNAiMAX according to the manufacturer’s protocol the following day. After 48hour of transfection, the transfected cells were trypsinized, stained with Trypan blue and counted using a hemocytometer. For the spheroid formation assay, 3000 cells were seeded in low attachment plates and spheroid formation was monitored by imaging via an inverted microscope at regular intervals.

### 2.10 Prediction of the Common Target Genes of the miRNA Pairs

The target genes of the miRNAs were predicted bioinformatically using TargetScan Human 7.1 (Agarwal *et al*., 2015) software. A cutoff of cumulative weighted context ++ score ≤-0.2 was used for selection of the target genes. The common targets of the miRNA pairs were selected, and the miRNA binding sites on the 3’-UTR sequence of the common target genes were predicted. The distances between the miRNA binding sites on the 3’-UTRs of the common target genes were calculated and checked using the UCSC genome browser. The common target genes that were found to be significantly upregulated in the TCGA-HNSC RNA-sequencing database (log (FC) ≥1, log(CPM) ≥2 and an FDR ≤0.05) and the EMBL Cancer Atlas database were selected for further downstream analysis.

### 2.11 Dose Kinetics Study

HEK-293T cells were seeded in 24-well plates to reach 60-70% confluence. On the day of transfection, the medium was replaced with antibiotic-free medium and 10ng of the recombinant pmirGLO plasmid was transfected into the cells using Lipofectamine 2000. For plasmid transfection; 10ng, 25ng, 50ng, 75ng and 100ng of the recombinant pRNA-U6 plasmid was cotransfected and luciferase activity was measured after 48hour of transfection. For mimic transfection, 24hour after the transfection of recombinant pmirGLO plasmids, the media was replaced with antibiotic-free media, and 10pmol, 15pmol, 20pmol, 25pmol and 30pmol miRNA mimic were transfected into the cells using Lipofectamine RNAiMAX. Luciferase activity was measured 24hour after mimic transfection. A miRNA dose *versus* luciferase activity curve was constructed to determine the optimal dose required for synergistic effect.

### 2.12 Luciferase Assay

Ten nanograms of the recombinant pmirGLO plasmid was transfected into HEK-293T cells. Fifty nanograms of pRNA-U6 recombinant plasmid was cotransfected with 1 µL of Lipofectamine 2000 or 20pmol miRNA mimics was transfected after 24hour with 1 µL of Lipofectamine RNAiMAX. Luciferase activity was measured after 48hour of transfection using a Dual-Luciferase Reporter Assay System (Promega). On the day of assay, media was discarded, and 100 µL of 1X Passive Lysis Buffer was added to each well and incubated at RT for 5 min. The cell lysate was collected in a prechilled 1.5 mL centrifuge tube, subjected to centrifugation at 13,000 rpm for 5 min at 4°C and 20 µL of the supernatant was collected in a fresh pre-chilled 1.5 mL centrifuge tube placed on ice. Hundred microliter of LAR II substrate solution (reconstituted with LAR II Buffer) and 100 µL of Stop & Glo solution (1000 µL of Stop & Glo Buffer + 20 µL of Stop & Glo substrate) were added to the samples to measure firefly luciferase (inducible) and Renilla luciferase (control) activities respectively using Glomax 20/20 luminometer (Promega). The luciferase activities were normalized to that of the transfection control (Renilla luciferase activity), and the activity of the target genes following miRNA overexpression was estimated as the relative luciferase activity (RLA) compared with that of the negative control.

### 2.13 Construction of Insertion/Deletion Plasmids

The NEBuilder Assembly Tool (version 2.7.1; NEB) was used for primer design. Primers flanking the miRNA binding sites on the 3’-UTR constructs were designed for amplification of the deletion constructs by inverse PCR. The primers for the spacer sequences used for insertion were designed from the 3’-UTR sequence of the CCNJ gene. The targets of the spacer sequences were also checked bioinformatically. Inverse PCR of the 3’-UTR constructs was performed using the KAPA LongRange HotStart PCT Kit (Sigma) following the manufacturer’s protocol, and the PCR products were purified using a QIAquick PCR Purification Kit (Qiagen). The spacer sequences were amplified from genomic DNA (3’-UTR of the CCNJ gene) using Taq polymerase according to the manufacturer’s protocol. The assembly of the insertion and deletion constructs was performed using NEBuilder® HiFi DNA Assembly Master Mix (NEB). For deletion, 50 ng of the amplified linearized vector was mixed with 10 µL of NEBuilder HiFi DNA Assembly Master Mix, and the volume was adjusted to 20 µL with ddH2O. For insertion, 50 ng of the amplified linearized vector was mixed with 2-fold molar excess of each insert, and 10 µL of NEBuilder HiFi DNA Assembly Master Mix, and the volume was adjusted to 20 µL with ddH2O. The samples were incubated in a thermocycler at 50°C for 60 min followed by an infinite hold at 4°C. Two microliters of the chilled assembled product were added to 100 µL of *E. coli* dH5α competent cells, after which transformation was performed. On the following day, positive colonies were selected by colony PCR with vector specific primers.

### 2.14 Gene Expression Study

The cells were seeded and allowed to reach 70-80% confluence within 24hour. The following day, the media was replaced with antibiotic free media and the cells were transfected with 20pmol miRNA mimic/s using Lipofectamine RNAiMAX. Total RNA was isolated from the transfected cells 48hour after transfection using RNeasy Mini Kit (Qiagen). One microgram of RNA was used for cDNA synthesis using Verso cDNA Synthesis Kit (Thermo) according to the manufacturer’s protocol (AB-1453-A, Verso cDNA Synthesis Kit, rev8). Ten nanograms of cDNA was mixed with 2X iTaq™ Universal SYBR® Green Supermix (Bio-Rad) and 300nmoles of gene specific forward and reverse primers in a total volume of 10µL to check the expression of the target genes by qRT-PCR. The expression of the common and unique target genes in the mimic transfected cells was normalized with respect to that in the negative control cell-line.

### 2.15 Protein Isolation and Western Blotting

Total protein was isolated from the transfected cells after 48hour of transfection using the SDS lysis method. The cells were trypsinized, and washed and 300μL of SDS lysis buffer supplemented with protease inhibitor cocktail was added to the cell pellet. The samples were incubated on ice for 30-40min, sonicated at 20% amplitude for 10min at 4°C using a Covaris M220 Focused Ultrasonicator (Covaris) and centrifuged at 12,000rpm at 4°C for 30min. The supernatant was collected, and the protein concentration was quantified using a PierceTM BCA Protein Assay Kit (Thermo Scientific) according to the manufacturer’s protocol (Pierce BCA Protein Assay Kit User Guide; Pub.No. MAN0011430 C00).

Proteins isolated from the cell-lysates (7.5-15μg) were subjected to denaturing 10% SDS-PAGE along with 10μL protein standard (Bio-Rad). Samples were electrophoresed using 1X Tris/Glycine/SDS Buffer at 100V for 60min. The proteins were transferred to a PVDF membrane using a Trans-Blot-Turbo System (Bio-Rad) at a constant voltage of 25V and 2.5A current for 3min. The membrane was blocked with 3% BSA for 2hour at RT, followed by overnight incubation with primary antibodies at 4°C. The membrane was washed with 1X TBST, and HRP-conjugated secondary antibodies were added. Afterwards, the membrane was incubated for 2 hours at RT and washed with 1X TBST. Afterwards, chemiluminescence-based detection of the bands was performed using Clarity ECL Western Substrate (Bio-Rad).

### 2.16 Proteomic assay

#### 2.16.1 In-sol trypsin digestion

A total of 100µg of protein sample was denatured with 8M urea, and the reaction mixture was diluted to 50µl with 50mM ammonium bicarbonate. Dithiothreitol (10mM) was used for reducing disulfide bonds for up to 1hour at 37°C followed by alkylation with 20mM iodoacetamide in the dark. After diluting the urea solution to less than 1M, the proteins were digested with trypsin at 1:50 overnight at 37oC. Following digestion, the resulting peptide mixture was acidified using formic acid, desalted, and vacuum concentrated.

#### 2.16.2 LC and SWATH-MS

LC-MS/MS was performed using an Eksigent microLC, connected on line with a TripleTOF 5600 mass spectrometer (Sciex). Desalted peptides were loaded in trap – elute mode with a reversed phase C18 column. The mobile phase for HPLC was as follows: water/acetonitrile/formic acid (A, 98/2/0.1%; B, 2/98/0.1%). Analytical separation was established by the following gradient procedure: initial 5% B for 5min, followed by a linear gradient from 3% B to 25% B for 68min, followed by another linear gradient to 35% B for 73min. Following the peptide elution window, the gradient was increased to 80% B in 2min and held for 3min. The initial chromatographic conditions were restored for 1min and maintained for 8min. SWATH-MS data were acquired using an ESI ion source with a voltage of 2300V, curtain gas of 25psi, nebulizer gas of 20psi, heater gas of 10psi, and source temperature of 130°C. Using different isolation widths, a set of 83 overlapping windows was constructed covering the mass range 350-1250 Da. The collision energy for each window was determined based on the appropriate collision energy set automatically with a spread of 5eV. The total duty cycle was 4.02 seconds.

*Data processing and analysis:* Identification of proteins was performed via database searching against the UniProt *Homo sapiens* database (June 2023), with DIA-NN (version 1.8.1) to obtain the in-silico spectral library. The search parameters were as follows: precursor FDR 1%; mass accuracy set to 50Lppm, scan window set to 0, isotopologues and MBR turned on, protein inference at the gene level, heuristic protein inference enabled, quantification strategy set to Robust LC (high precision), neural network classifier single-pass mode, and cross-run normalization RT dependent. In library-free mode, the main search settings were the same with additional settings for in silico library generation as follows: Trypsin/P with maximum 2 missed cleavage; protein N-terminal M excision on; Carbamidomethyl on C as a fixed modification; no variable modification; peptide length from 7 to 30; precursor charge 1–4; precursor m/z from 350 to 1250; and fragment m/z from 100 to 1500.

### 2.17 Metabolomic assay

#### 2.17.1 Metabolite extraction and derivatization

Cells were seeded to reach 70-80% confluence within 24hour. The following day, the media was replaced with antibiotic free media, and the cells were transfected with 20pmol miRNA mimic/s using Lipofectamine RNAiMAX. On the day of the assay, the cells were washed twice with 1ml of 150mM NaCl. Then, 500µl of prechilled extraction solvent comprising of water: isopropanol: acetonitrile (2:3:3) containing an internal standard (10 µM 3-amino 4-methoxy benzoic acid, aka, 3AMBA) was added to the plate, and the cells were collected using a cell scraper, stored on ice and processed immediately. The samples were further subjected to three freeze-thaw cycles using liquid nitrogen and centrifuged at20000 x g for 30mins at 4°C. A total of 150µl of supernatant was transferred to a GC vial and evaporated to dryness under vacuum. Dried samples were then derivatized with 30µl MOX at 50°C for 1hour followed by 50µl of MSTFA at 65°C for 1hour. Pooled QC and extraction blank samples were also prepared in a similar manner and analyzed using GCMS. Along with these, QC samples were also prepared by pooling 50µl of supernatant from each sample. Pooled QC and extraction blank samples were also derivatized as described above.

#### 2.17.2 GCMS analyses

Derivatized samples were analyzed with a 7890B GC fitted with a HP-5MS UI column (30 m × 0.25 mm × 0.25μm) coupled to a 5977B single-quadrupole mass spectrometer (Agilent, USA) with helium as the carrier gas. One microliter of each sample was injected into the front inlet maintained at 300°C in split-less mode in a randomized manner with intermittent injection of pooled QC and blank samples. Initially, the oven temperature was maintained at 70°C for 5min, followed by a gradual increase at 5°C/minute up to 280°C. The temperature was further increased to 295°C at 10°C/minute and held at 295°C for 4min. MS source and MS quad temperatures were set to 230°C and 150°C, respectively. The mass spectra were acquired in full scan mode in the m/z range of 45-600.

#### 2.17.3 Metabolomic data analysis

Following manual inspection of chromatographic performance and instrument response, MassHunter quantitative software (Agilent,USA) was used for data deconvolution, feature extraction and integration. Typically, one quantifier and two qualifier ions were used to extract features of interest. Putative compound identification was based on matching with the NIST 14 library as well as comparing the retention time and fragmentation with those of authentic standards wherever available. The data table comprising the area under the curve for individual features for the respective samples was used to calculate the coefficient of variation (CV) of the respective internal standard-normalized features. Only those features with a CV < 30% in pooled QC samples were used for further univariate and multivariate analysis. Sum-normalized, log-transformed and Pareto-scaled data were used for multivariate analysis via MetaboAnalyst 5.0 (https://www.metaboanalyst.ca/). Unsupervised principal component analysis (PCA) was used to check the quality of the data as well as any overall divergence of metabotypes among the control and treatment groups. Partial least square discriminant analysis (PLS-DA) was used for supervised analysis to test the utility of the metabotype in class discrimination (between control and miRNA-treated samples). Heatmap analysis was used to analyze the clustering pattern of control vs treated samples. Volcano plot analysis was used to identify features that showed significant changes in abundance (fold-change > 1.5) upon miRNA treatment after FDR-correction. The statistical significance of changes in the sum-normalized relative abundance of features of interest (with respect to the control group) was further tested using ANOVA with the Dunnet test in GraphPad Prism 9(GraphPad Software, Boston). P < 0.05 was considered to indicate statistical significance.

### 2.18 Statistical Analysis

The significance of the deregulation of miRNAs in paired tissue samples was determined by paired t-test. A p-value less than 0.05 was considered to be significantly differential among normal and disease samples. The significance of the differentially expressed miRNAs or genes in cell-lines was determined using Mann-Whitney U test. The p-values were calculated using GraphPad (PRISM) software (GraphPad Prism version 5.0.0 for Windows, GraphPad Software, San Diego, California USA, www.graphpad.com). KEGG pathway analysis was performed using ShinyGO 7.7 (http://bioinformatics.sdstate.edu/go77/), and an FDR cutoff <0.05 was used for the selection of the top enriched pathways.

## 3. RESULTS

### 3.1 Identification and Validation of the Common Deregulated miRNAs in Oral Cancer

To identify the deregulated tumor-suppressor miRNAs, we conducted genome-wide small RNA sequencing using four paired OSCC and adjacent normal tissues, obtained from the oral cancer patients from Eastern India. We identified 57 differentially regulated miRNAs including 23 upregulated and 34 downregulated miRNAs in OSCC after applying the criteria of |Log FC|≥1, Log(CPM)≥2 and an adjusted P-value≤0.05 (**Table S4**) (**Figure 1a**). We compared our small RNA-seq data with that of 303 HNSCC and 32 normal tissue samples from The Cancer Genome Atlas (TCGA) for identification of common deregulated miRNAs. Approximately, 84% of the differentially expressed miRNAs (41 miRNAs) in our cohort showed similar deregulation with the TCGA-HNSCC cohort, suggesting their universal regulation in OSCC development (**Figure 1b**). To further validate our data, 13 common deregulated miRNAs were randomly selected and validated in an additional 33 paired OSCC and adjacent normal tissue samples using qRT-PCR. The validation data showed ∼85% concordance with the small RNA-seq data (**Figure S1**).

**Figure 1:**
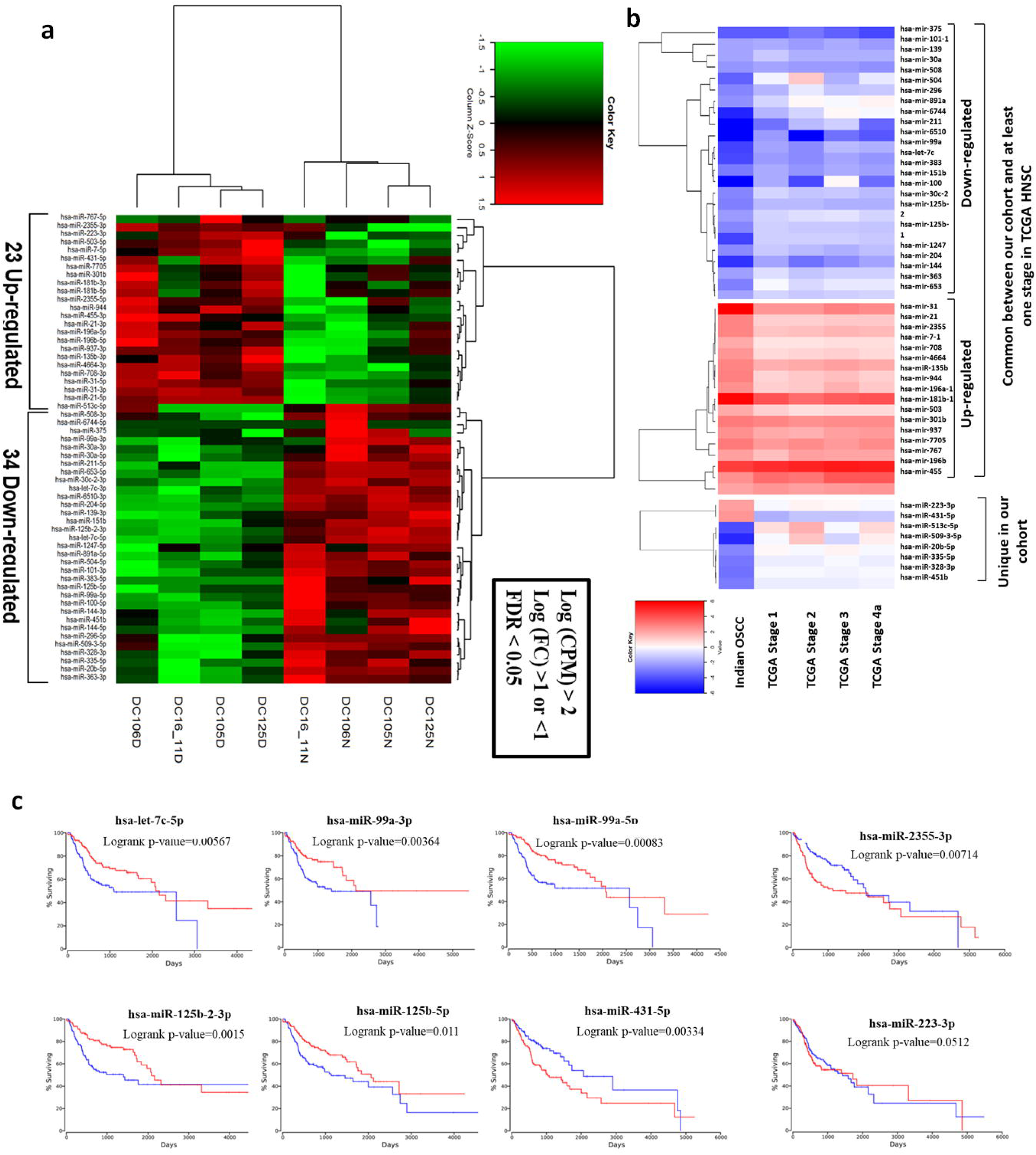
Identification of the clinically significant deregulated miRNAs in oral cancer. **a)** Heatmap representing the 57 deregulated miRNAs in OSCC *versus* adjacent normal tissue (23 up-regulated and 34 down-regulated miRNAs) identified from the small RNA-sequencing having a |Log FC|≥1, Log cpm≥2, FDR≤0.05). **b)** Heatmap representing the 41 common and 8 uniquely deregulated miRNAs in OSCC tissue compared to normal between the small RNA-sequencing data and TCGA-HNSC small RNA-sequencing database. **c)** Kaplan-Meier survival plots of the 8 common deregulated miRNAs showing significant correlation with disease prognosis (Mantel-Cox log-rank p-value≤0.05).

### 3.2 Identification of the Clinically Significant Tumor Suppressor miRNAs in Oral Cancer

We used Kaplan-Meier survival analysis on all 41 deregulated miRNAs in order to determine which ones were clinically significant. Of these, eight miRNAs, seven of which were downregulated and one of which was upregulated, exhibited noteworthy correlations and prognostic significance among patients with HNSCC. Patients with HNSCC had poor survivability when hsa-let-7c-5p, hsa-miR-99a-3p, hsa-miR-99a-5p, hsa-miR-125b-2-3p, and hsa-miR-125b-5p were downregulated and hsa-miR-2355-3p was upregulated. However, patients with low expression of the downregulated miRNAs hsa-miR-204-5p and hsa-miR-100-5p had significantly higher survivability than those with high expression (**Figure 1c**). Additionally, miR-204-5p, let-7c-5p, miR-99a-5p, and miR-125b-5p were significantly downregulated in OSCC and widely expressed in all samples (Table S5). The nonspecific amplification of miR-99a-5p may be the cause of the three miRNAs, miR-99a-5p, miR-99a-3p, and miR-100-5p, which are all members of the same miRNA family. Thus, for additional downstream research, we have chosen three down-regulated miRNAs: miR-204-5p, let-7c-5p, and miR-125b-5p. The ECM-receptor interaction, MAPK signaling, JAK-STAT signaling, PI3K-Akt signaling, and p53 signaling are among the key pathways that are dysregulated in cancer, according to KEGG pathway analysis using the target genes of these three miRNAs (Table S6). The majority of these pathways, such as the p53 and MAPK signaling pathways, are implicated in the control of cancer cell division and proliferation [29, 30].

In order to the role of these miRNAs in OSCC, ORL-48 oral cancer cells were transduced with lentivirus particles for constitutive overexpression of these 3 miRNAs (**Figure S2a**). For the purpose of functionally characterizing the miRNAs, these cell lines were put through phenotypic and biochemical assays in comparison to the scramble control cell line. Cellular proliferation in ORL48 cells was lower when miR-125b-5p, let-7c-5p, and miR-204-5p were overexpressed than in the scramble control cell line (**Figure S2b**). Overexpression of miR-204-5p also decreased the rate of wound closure, according to a scratch wound-healing cell migration assay (**Figure S2c**). On the other hand, cellular apoptosis was not substantially altered by the overexpression of any of the three miRNAs (**Figure S2d**). These results demonstrated the involvement of these miRNAs in the regulation of cell proliferation and migration in OSCC.

### 3.3 Identification and Validation of Common Targets of miR-125b-5p, let-7c-5p and miR-204-5p

We predicted 927, 1191 and 788 target genes of miR-125b-5p, let-7c-5p and miR-204-5p, respectively. Cytotoxicity is frequently the result of administering a single miRNA for effective treatment. Therefore, we investigated the cooperative effect of miRNA pairs. While, 182, 130 and 118 genes were targeted by miR-125b-5p and let-7c-5p pair, let-7c-5p and miR-204-5p pair, and miR-125b-5p and miR-204-5p pair, respectively. Since these 3 miRNAs were downregulated in OSCC, the upregulated common target genes of these miRNA pairs in oral cancer were identified from the TCGA-HNSC RNA-sequencing data using cutoff values of Log(FC) ≥1, Log(CPM) ≥2 and FDR< 0.05. We identified 34, 11 and 7 common target genes for miR-125b-5p and let-7c-5p, let-7c-5p and miR-204-5p and miR-125b-5p and miR-204-5p, respectively (**Figure S3a**). The pre-miRNA sequences of the three downregulated miRNAs were cloned into the pRNA-U6 vector, and the 3’-UTR sequences of 28 randomly chosen common target genes were cloned into the pmirGLO vector for validation. HEK-293T cells were co-transfected with an optimal dosage of 10ng of the recombinant pmirGLO plasmid and 50ng of the recombinant pRNA-U6 plasmid/s (**Figure S3b**). Luciferase activity was assessed 48 hours after transfection. Following treatment with either let-7c-5p or miR-125b-5p, the 20 genes that shared target sites with both proteins exhibited noticeably lower luciferase activity. Likewise, two genes with common target sites for miR-125b-5p and miR-204-5p, as well as all five genes with common target sites for let-7c-5p and miR-204-5p, demonstrated markedly decreased luciferase activity by either of these miRNAs, confirming the efficient translational repression of the selected genes by these 3 miRNAs (**Figure S3c**).

### 3.4 The Synergistic Effect of miRNAs

To study the synergistic effect of miRNA-pairs, cells were co-transfected either with 50ng of individual miRNA-overexpressing plasmids or with 25ng of pair of miRNA-overexpressing plasmids. A significant reduction in luciferase activity was observed after co-transfection with miRNA-pairs compared to the geometric mean of the luciferase activities with individual miRNAs, indicating a synergistic downregulation of their luciferase activity (**Figure S3d**). We conducted a more controlled experiment and investigated the synergistic effect of miRNAs by transfecting individual miRNA mimics or co-transfecting miRNA mimic pairs in order to corroborate our findings. A dose kinetic study using increasing doses of miRNA mimics (0pmol, 10pmol, 15pmol, 20pmol, 25pmol, and 30pmol) was used to optimize the dose. For downstream experiments, a standard dose of 20 pmol was determined to be optimal (**Figure S4a-b**). When compared to the geometric mean of the luciferase activities with individual miRNA mimics for the genes HK2, BCAT1, NKIRAS2, CCDC71L, USP38, MTUS1, LBH, LRRC10B, BTG2, PTAR1, C15ORF39, PTPRD, HAS2, HMGA2, ACER2, TET3, MIB1, and ERCC6, co-transfection with half the doses of miRNA mimic-pairs (10 pmol each) significantly decreased the luciferase activity. There was no discernible decrease in luciferase activity for the remaining 10 target genes (**Figure 2a**).

**Figure 2:**
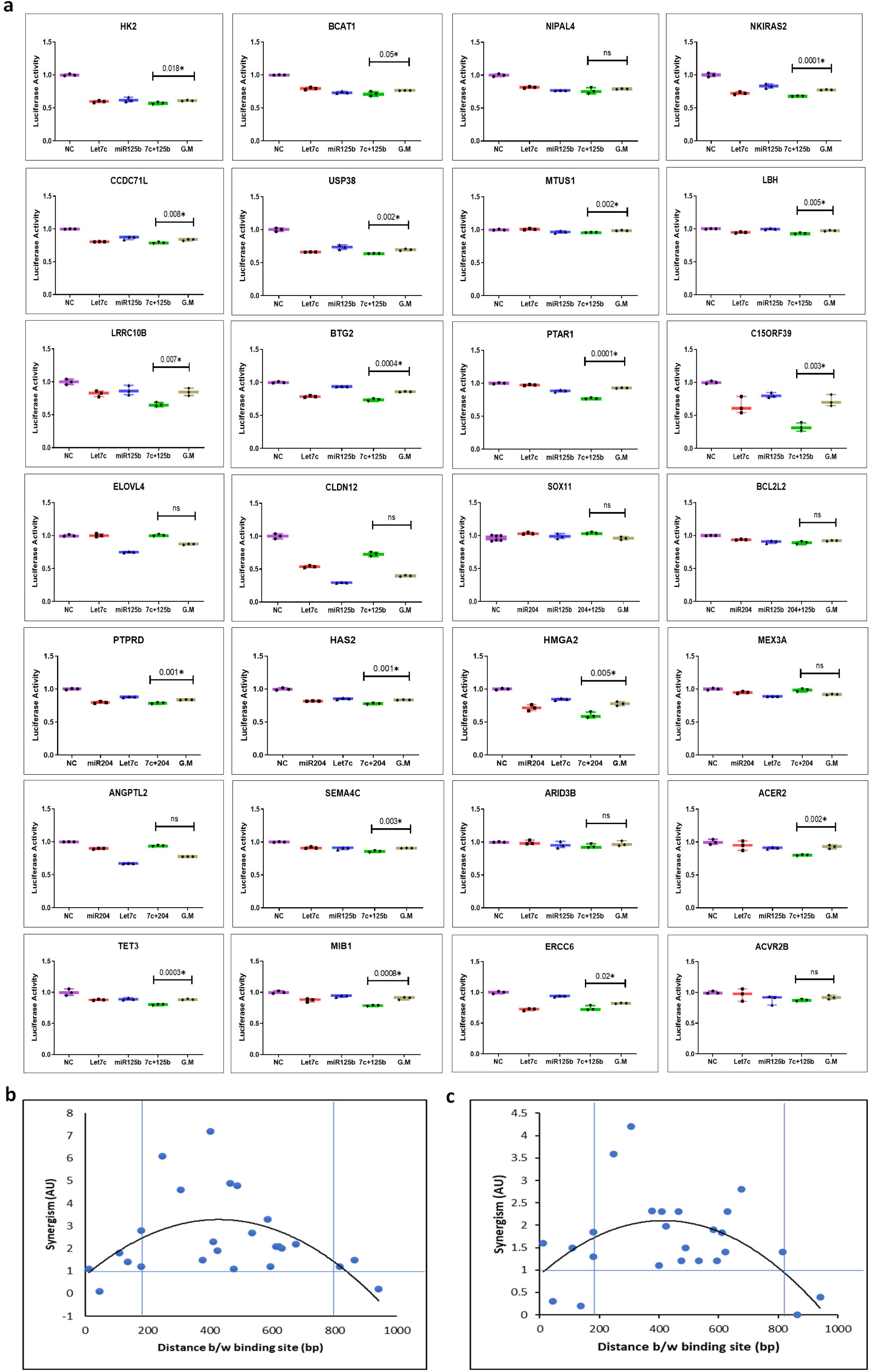
MiRNA-mediated synergistic down-regulation of the common target genes. **a)** Luciferase assay representing the synergistic down-regulation of the common target genes upon co-expression of half the dose (10pmol) of miRNA mimics in HEK-293T cells. **b)** Distance between MRE of miRNA-pairs *versus* synergism curve with co-expression of half the dose (25ng) of recombinant pRNA-U6 plasmids. **c)** Distance between MRE of miRNA-pairs *versus* synergism curve with co-expression of half the dose (10pmol) of miRNA mimics. G.M: Geometric Mean; p*≤0.05, ns>0.05.

The miRNA-mediated synergism was found to be significantly influenced by the distance between the MREs of the common target genes. Distances between the two MREs ranged from 180 bp to 816 bp for the 18 target genes that demonstrated synergistic decreases in luciferase activity with co-expression of the miRNA-pairs, while the remaining 10 genes had a distance beyond that range. To further validate our findings, we determined the percentage reduction in luciferase activity and calculated the effect of synergism (*E_syn_*) (**Table S7**) with the following equation:

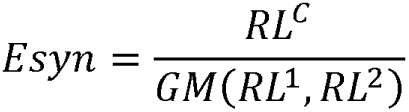

where, *RL^C^*= the percent reduction in luciferase activity associated with co-expression of 10pmol of each miRNA mimics or 25ng of recombinant pRNA-U6 plasmid, *RL^1^*= the percent reduction in luciferase activity with expression of 20pmol of mimic 1 or 50ng of recombinant pRNA-U6 plasmid 1, *RL^2^*= the percent reduction in luciferase activity with expression of 20pmol of mimic 2 or 50ng of recombinant pRNA-U6 plasmid 2, and G.M= the geometric mean of *RL^1^* and RL^2^

The effect of synergism was defined as follows: (i) *E_syn_*>1: Synergistic Effect, (ii) *E_syn_* =1: Additive Effect, and (iii) *E_syn_* <1: Antagonistic Effect. For every target gene of the miRNA mimic pairs (**Figure 2c**) or recombinant pRNA-U6 plasmid pairs (**Figure 2b**), we plotted the distance between the MREs and *Esyn*. An optimal distance of 200 to 800bp was found to show synergistic effects with co-transfection by the miRNA-pairs compared to the geometric mean of individual miRNAs (**Figure 2c-d**).

### 3.5 The Role of the Distance between MRE in Synergism

To further elucidate the role of distance between MREs, we selected the HK2, BCAT1 and CCDC71L pmirGLO-3’-UTR plasmids, and generated 9 insertion/deletion constructs having different distances between the MREs ranging from 101bp to 1090bp. The plasmids were co-transfected either with 50ng of pRNA-U6 recombinant plasmids individually or in combination of 25ng of each miRNA pairs (**Figure 3a**). The distance versus synergism curve for the insertion/deletion constructs also showed that an optimal distance of 200-800bp between the MREs of their common target genes had synergistic effects (**Figure 3b**) (**Table S8**). Similar results were obtained when we used 20pmol of individual miRNA mimics or a combination of 10pmol of each miRNA mimic pair (**Figure 3c-d**) (**Table S8**).

**Figure 3:**
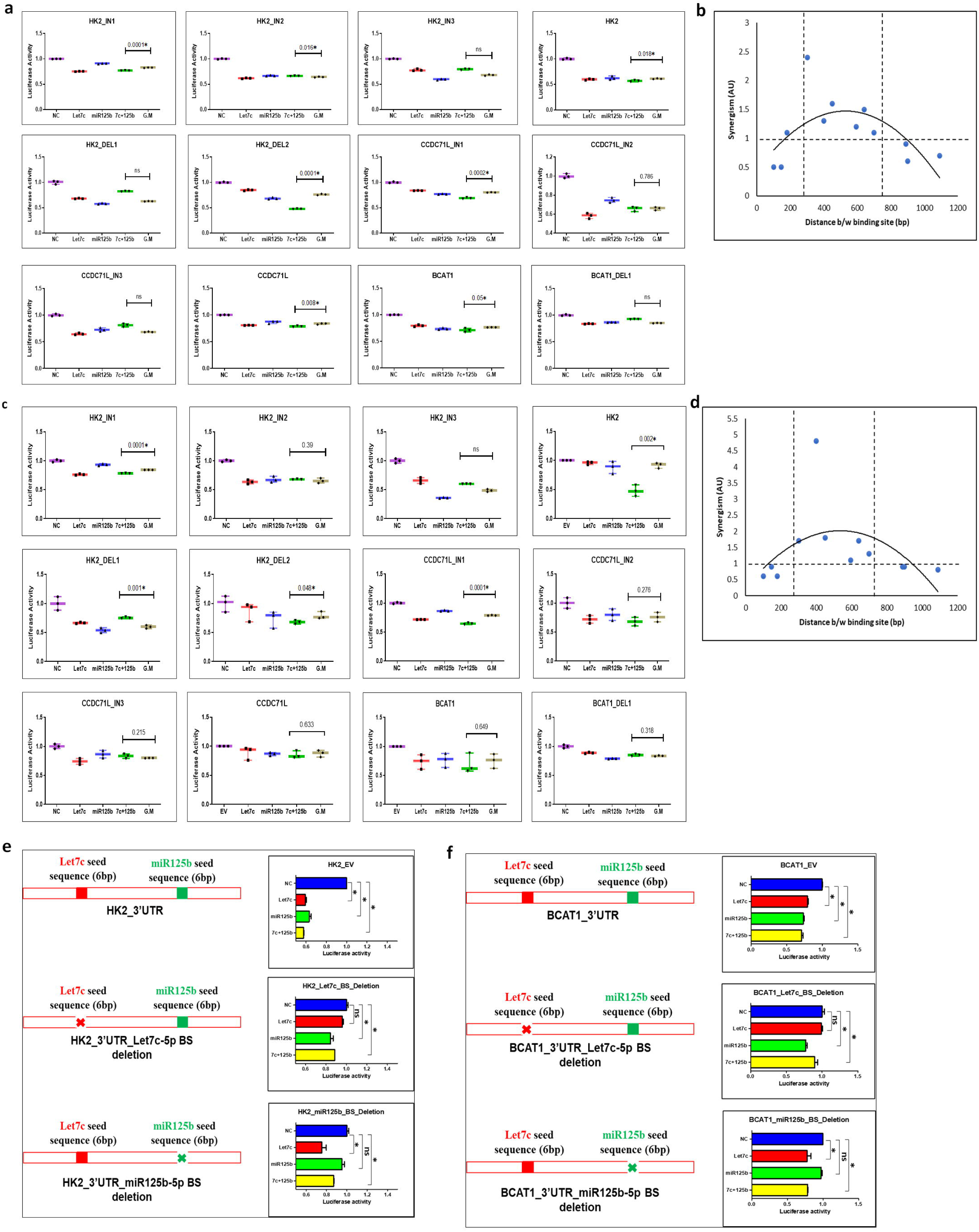
The role of distance between MRE of miRNA-pairs on the 3’-UTR of their common target genes on synergism. **a)** Luciferase assay representing the synergistic down-regulation of the insertion/deletion constructs with co-expression of half the dose (25ng) of recombinant pRNA-U6 plasmids **b)** Distance between MRE of miRNA-pairs *versus* synergism curve with co-expression of half the dose (25ng) of recombinant pRNA-U6 plasmids **c)** Luciferase assay representing the synergistic down-regulation of the insertion/deletion constructs with co-expression of half the dose (10pmol) of miRNA mimics **d)** Distance between MRE of miRNA-pairs *versus* synergism curve with co-expression of half the dose (10pmol) of miRNA mimics using insertion/deletion 3’-UTR constructs. **e-f)** Luciferase assay representing the deletion of either let-7c-5p or miR-125b-5p seed sequences from the 3’-UTR of HK2 and BCAT1 constructs resulting in a loss of synergism. DEL: Deletion; IN: Insertion; BS: Binding site; p*≤0.05, ns>0.05.

To further verify whether the synergistic effect is actually mediated by miRNA binding to the MREs, the 6bp seed sequences of let-7c-5p or miR-125b-5p was deleted from the MREs of the 3’-UTR of HK2 and BCAT1 genes. Deletion of either of the seed sequences from the 3’-UTR constructs did not show any significant change in the luciferase activity for the specific miRNA (**Figure 3e-f**). Moreover, deletion of any of the seed sequences resulted in a loss of synergistic effect by the miRNA mimic-pairs (**Figure 3e-f**).

### 3.6 MiRNA-mediated Synergism Amplifies Target Specificity and Reduces Off-targets

To investigate the effect of miRNA-mediated synergism in oral cancer, we selected two tumor-suppressor miRNAs, miR-125b-5p and let-7c-5p, which are downregulated in OSCC. MiRNA mimics were transfected into the oral cancer cells either individually (20 pmol) or in combination (10 pmol each). The expression of both the common and unique target genes were found to be downregulated in individual miRNA mimic transfected cells compared to the negative control cells (**Figure 4a-b**). Moreover, the common target genes, such as HK2, BCAT1, NKIRAS2, and CCDC71L exhibited synergistic downregulation of gene expression following co-transfection of half doses of the miRNA mimic-pair, as also observed in our luciferase assay (**Figure 4a**). In contrast, the unique target genes did not show any significant synergistic downregulation of their target genes (**Figure 4b**). These observations suggest that miRNA mediated synergism might result in effective downregulation of common target genes while reducing off-target effects. The protein expressions of HK2 and BCAT1 were also determined following the transfection of individual miRNA mimics or co-transfection of miRNA mimic-pairs with half dose of each miRNA. We observed 5% and 1% reduction in HK2 protein expression with transfection of let-7c-5p and miR-125b-5p respectively, while 9% and 0% reduction in the BCAT1 protein were observed following the transfection of respective miRNAs. Co-transfection of the miRNA mimic-pairs in the cancer cells resulted in 65% and 13% reductions in HK2 and BCAT1 protein expressions, respectively, further validating the synergistic effect of miRNAs in regulating their target genes (**Figure 4c**).

**Figure 4:**
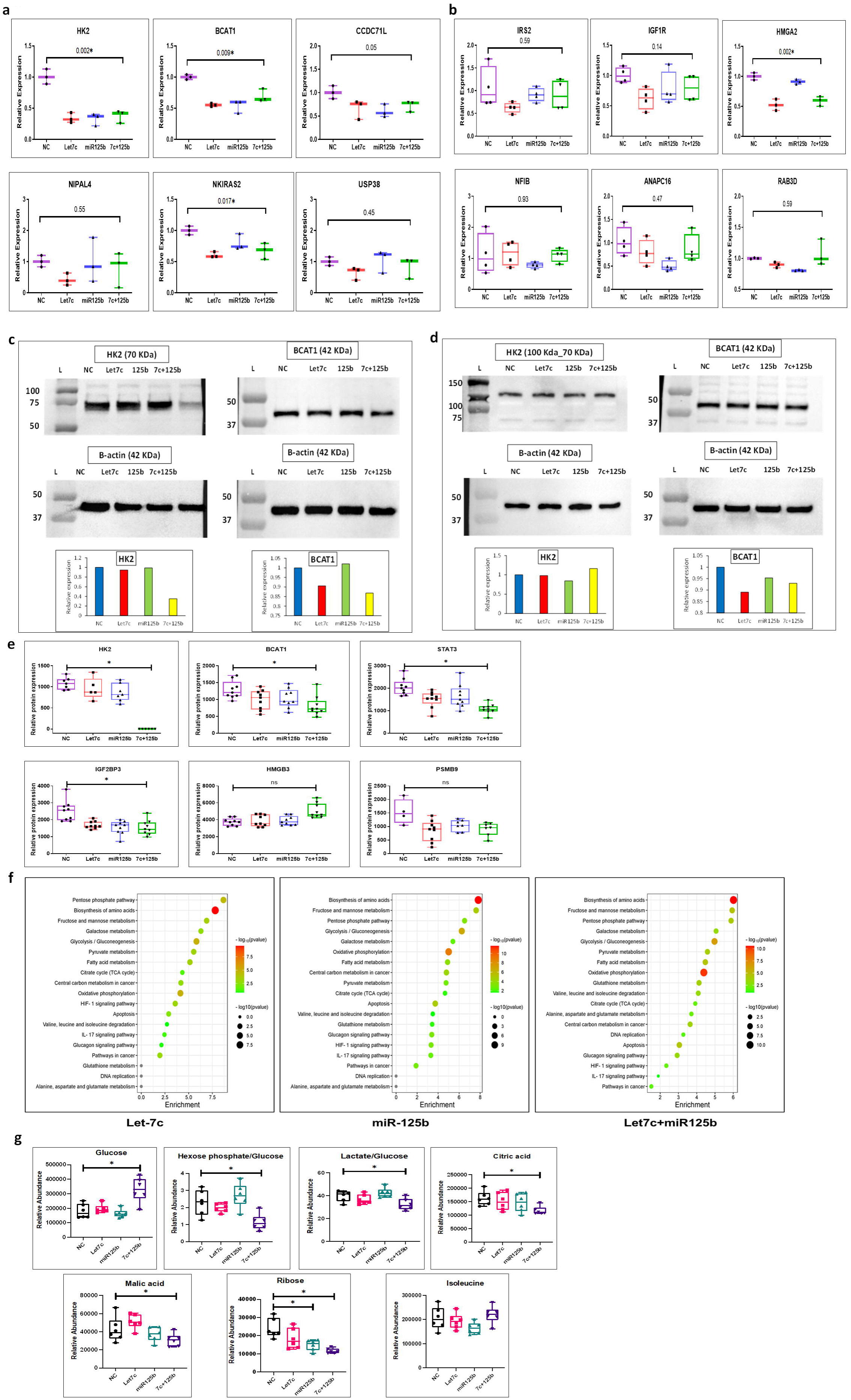
The effect of miRNA-mediated synergism on off-targets and cellular toxicity. **a)** Gene expression of the common target genes of let-7c-5p and miR-125b-5p (HK2, BCAT1, CCDC71L, NIPAL4, NKIRAS2 and USP38) with co-expression of half the dose (10pmol) of miRNA mimics in ORL-48 cell-line. **b)** Gene expression of the unique target genes of let-7c-5p (IRS2, IGF1R and HMGA2) and miR-125b-5p (NFIB, ANAPC16 and RAB3D) with co-expression of half the dose (10pmol) of miRNA mimics in ORL-48 cell-line. **c)** Protein expression of the common target genes of let-7c-5p and miR-124b-5p (HK2 and BCAT1) with co-expression of half the dose (10pmol) of miRNA mimics in ORL-48 cell-line. **d)** Protein expression of the common target genes of let-7c-5p and miR-124b-5p (HK2 and BCAT1) with co-expression of half the dose (10pmol) of miRNA mimics in HaCaT cell-line. **e)** Protein expression of the common target genes (HK2, BCAT1, STAT3, AGO2) and the unique target genes of Let-7c-5p (IGF2BP3, CALU) and miR-125b-5p (HMGB3, PSMB9) (iTRAQ proteomics). **f)** Enrichment bubble-plot representing the top enriched significant pathways targeted by Let-7c-5p, miR-125b-5p and with co-expression of the miRNA-pairs (iTRAQ proteomics). **g)** Metabolomic assay representing the relative abundance of glucose-phosphate and lactate to glucose and the overall abundance of glucose and glucose-phosphate in ORL-48 cells upon co-expression of half the dose (10pmol) of miRNA mimics. L: Ladder; NC: Negative control mimic; p*≤0.05, ns>0.05. S1: ORL48 cells transfected with 20pmol NC mimic; S2: ORL48 cells transfected with 20pmol Let-7c-5p mimic; S3: ORL48 cells transfected with 20pmol miR-125b-5p mimic; S4: ORL48 cells transfected with 10pmol Let-7c-5p mimic and 10pmol miR-125b-5p mimic

### 3.7 The Role of miRNA-mediated Synergism in the Global Proteomic Profile of Oral Cancer Cells

To investigate the effect of miRNA-mediated synergism, we performed global protein expression profiling of oral cancer cells (ORL-48) following treatment of the individual miRNA mimic or miRNA mimic-pairs and compared the results to those of mock-treated cells using iTRAQ-based proteomic analysis. We first evaluated the protein expression of the unique and common target genes of let-7c-5p and miR-125b-5p following treatment with individual miRNAs or miRNA mimic-pairs. Among the 2053 proteins identified from the proteomics data, 6 proteins were found to be common targets of the miRNA-pair, while 24 and 19 proteins were uniquely targeted by let-7c-5p and miR-125b-5p, respectively (**Table S9**). Among the 6 common targets, 3 were upregulated in oral cancer cells, namely, HK2, BCAT1 and STAT3 and a significant reduction in their protein expression was observed in individual miRNA transfected cells. Moreover, we observed synergistic reductions of these common proteins following co-transfection of miRNA mimic-pairs (**Figure 4e**). The distances between the MREs of the miRNA-pairs on the 3’-UTR of HK2, BCAT1, and STAT3 were 401bp, 594bp, and 1448bp, respectively. Among the 24 and 19 unique target proteins of let-7c-5p and miR-125b-5p, 3 unique target proteins of let-7c-5p, namely, IGF2BP3, GALNT2, and CALU and 2 unique target proteins, HMGB3, and PSMB9 of miR-125b-5p, were found to be upregulated in oral cancer. The unique target genes of the respective miRNAs upregulated in oral cancer did not show any significant downregulation in their protein expression with co-expression of the miRNAs, indicating target specificity and reduction of off-target effects (**Figure 4e**). Among the remaining proteins that were predicted to be uniquely targeted by the respective miRNAs, many did not show any target specific deregulation. Although some of the proteins were significantly downregulated in the respective miRNA transfected cells, all these proteins were also significantly downregulated by the other miRNAs, suggesting that these proteins were the results of miRNA-independent downstream effects.

Next, we identified the significantly enriched pathways by the deregulated proteins in individual and co-transfected miRNA mimics compared to mock controls. Pathway enrichment analysis showed reduced enrichment of HK2 mediated pathways, such as, glycolysis, the TCA cycle, oxidative phosphorylation and pentose phosphate pathway in cells co-expressing miRNA mimic-pairs compared to those in the individual miRNA transfected cells, indicating an overall reduction in glucose metabolism and energy production (**Figure 4f**). Similarly, downregulation of BCAT1 was reflected by decreased enrichment of amino acid degradation pathway and increased branched chain amino acids synthesis pathway in cells co-transfected with miRNA mimic-pairs. In contrast, the individual mimic transfected cells exhibited enrichment of BCAA degradation but no enrichment of BCAA synthesis pathway (**Figure 4f**). Decreased enrichment of HIF-1 signaling and IL-17 signaling pathways in the co-transfected cells compared to that of the individual mimic transfected cells indicated a reduction in inflammatory cytokines and cellular stress, in turn resulting in reduction of overall cellular toxicity (**Figure 4f, Table S10**).

### 3.8 The Role of miRNA-mediated Synergism in the Metabolomic Profile of Oral Cancer

Combined treatment of let-7c-5p and miR-125b-5p resulted in significant synergistic downregulation of HK2 and BCAT1. HK2 catalyses the first rate-determining step of glycolysis resulting in the conversion of glucose to glucose-6-phosphate. BCAT1 on the other hand facilitates the reversible transamination of branched-chain amino acids (BCAAs). To further validate the effect of synergistic downregulation of HK2, we performed metabolomics assays in individual or combined miR-125b-5p and let-7c-5p mimics transfected oral cancer cells (**Figure S4**). A significant reduction in glucose-phosphate/glucose, lactate/glucose ratio and an increase in glucose was observed with co-expression of half the dose of the miRNA-pairs, indicating an overall reduction in glycolysis. In contrast, no significant deregulation of these metabolites were observed in individual mimic transfected cells (**Figure 4g**). The metabolomic assay also indicated the effect of reduced BCAT1 expression. Transfection of the individual miRNAs showed a reduction in isoleucine levels in oral cancer cells, while co-expression of half doses of the miRNA-pair showed an marginal increase in the abundance of isoleucine (BCAA) compared to the individual mimic transfected cells (**Figure 4g**). Since, downregulation of BCAT1 resulted in enrichment of the BCAA synthesis pathway in the co-transfected cells from the proteomics data, an increase in isoleucine further validated the synergistic effect of miRNAs in our metabolomics analysis.

### 3.9 The Synergistic Role of miRNAs in Oral Cancer Phenotype

To study the synergistic effect of miRNAs, oral cancer cells were either transfected with 20pmol of the individual miRNA mimics or co-transfected with 10pmol (one-half of the optimal dose) of each of the miRNA mimics followed by functional characterization. We performed WST1 cell proliferation and scratch wound-healing assays with the transfected cells. A reduction in cell proliferation was observed with overexpression of all 3 miRNAs individually as well as in combination. Moreover, co-transfection of a half dose of the miRNA-pairs showed significant reduction in cell proliferation compared to the individual mimic transfected cells, indicating a synergistic effect of the miRNA combination (**Figure 5b**). We observed a reduction in cell migration with overexpression of the individual miRNA mimics, while co-expression of half dose of miRNA mimic-pair resulted in a significantly higher reduction in cell migration. Overexpression of miR-204-5p individually or in combination with either of the two miRNA mimics showed a greater reduction in cell migration rate and synergism indicating the involvement of mir-204-5p in the regulation of cell migration in OSCC, consistent with our previous observations (**Figure 5a**). These findings indicate the probable role of these miRNAs in the synergistic regulation of cell proliferation and migration in oral cancer.

**Figure 5:**
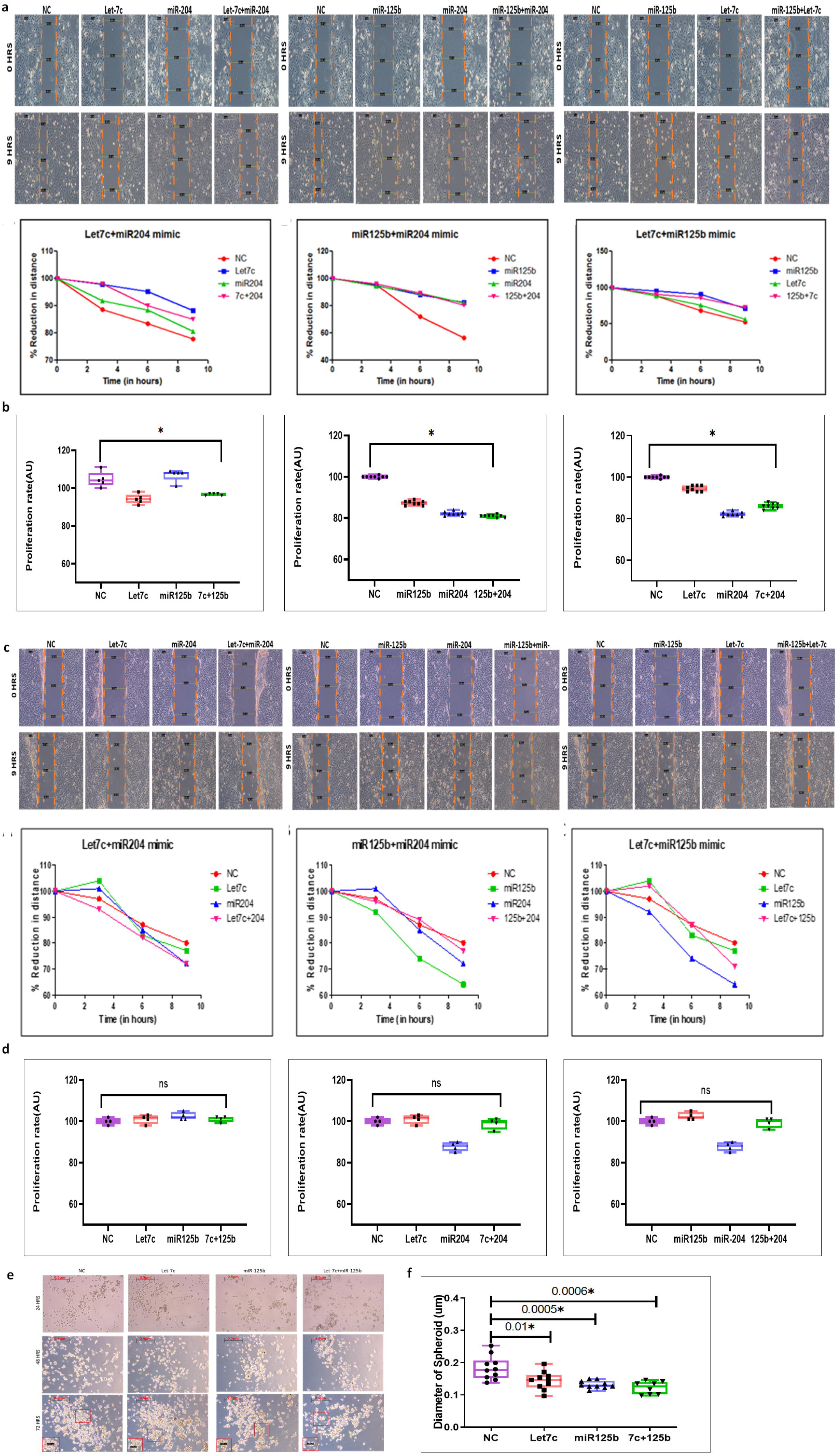
The role of miRNA-mediated synergism in oral cancer phenotype. **a)** Scratch wound-healing cell migration assay and **b)** WST1 cell proliferation assay with co-transfection of half the dose (10pmol) of miRNA mimics in ORL-48 cell-line. **c)** Scratch wound-healing cell migration assay and **d)** WST1 cell proliferation assay with co-transfection of half the dose (10pmol) of miRNA mimics in HaCaT cell-line. **e)** 3-D cancer spheroid formation assay with co-transfection of half the dose (10pmol) of miRNA mimics in ORL-48 cell-line. **f)** Representation of the reduction in spheroid diameter with co-transfection of miRNA mimics in ORL-48 cell-line. NC: Negative control mimic; p*≤0.05, ns>0.05.

### 3.10 Overexpression of miRNA pairs Synergistically Reduces Cancer Spheroid Formation

A synergistic effect was evident with co-expression of the miRNAs in the oral cancer cell-lines. However, *in-vitro* 3D cell culture models are widely recognized as more physiologically relevant systems compared to 2D formats. Hence, we studied the effect of miRNA co-expression in 3D cell culture models. Transfection of 20pmol of the individual mimics as well as co-transfection of one-half of the dose (10pmol) of the combination of miRNA mimics in ORL-48 cells showed a significant reduction in spheroid formation compared to the negative control cell-line (**Figure 5e-f**). Additionally, the co-transfected cells showed a greater reduction in the colony diameter of spheroids compared to that of the individual mimic transfected cells, indicating a synergistic reduction in cancer spheroid formation with co-transfection of the miRNA-pairs (**Figure 5e-f**).

### 3.11 MiRNA-mediated Synergism in Wildtype Cells and Their Effect on Cellular Toxicity

To study the effect of miRNA-mediated synergism in normal cells, individual miRNA mimics or half doses of miRNA mimic-pairs were transfected in a keratinocytes cell line (HaCaT). The relative protein expression of HK2 and BCAT1 was quantified in mock treated, individual miRNA or miRNA mimic-pair transfected HaCaT cells. 2% and 16% reduction in HK2 and 11% and 5% reduction in BCAT1 protein were observed with transfection of let-7c-5p and miR-125b-5p mimics, respectively, while a marginal reduction in the protein expression of BCAT1 (7%) and no reduction in HK2 (0%) protein were observed with co-transfection of half the dose of the miRNA mimic-pairs. The protein expression of both HK2 and BCAT1 was lower in normal cells compared to the oral cancer cells. No synergistic downregulation of the proteins was observed with co-expression of reduced doses of miRNA mimic-pair owing to their low abundance in normal cells (**Figure 4d**), suggesting level of gene expression is also another contributing factor to miRNA-mediated synergism. Transfection of the individual miRNA mimics or co-transfection of half dose of the miRNA-pairs in keratinocytes also did not significantly alter cellular proliferation (**Figure 5d**) or migration (**Figure 5c**). Hence, miRNA mediated synergism effectively reduces cell proliferation and migration of the cancer cells, but does not alter the phenotype of the normal keratinocytes.

## 4. DISCUSSION

MiRNAs play vital roles in tumor initiation, progression, and metastasis[31]. The ability of miRNAs to regulate important cellular processes by concurrently regulating multiple targets illustrates their potential as a viable therapeutic tool [32, 33]. One of the major drawbacks of miRNA therapeutics is the cellular toxicity, induced by off-target effects [4, 10]. Reduced doses of combinations of miRNAs can increase the specificity of their targets, in turn reducing cellular toxicity due to the co-operative nature of interactions between the miRNAs and the 3’-UTR of target genes. This strategy might serve as a novel therapeutic approach for cancer treatment.

Here, we identified the potential factors that might regulate miRNA-mediated synergism, and applied this approach to investigate the effect of miRNA-mediated synergism in oral cancer. To identify potential tumor suppressor miRNAs, genome-wide small RNA-sequencing data from OSCC patients from Eastern India were compared with TCGA-HNSCC data. Among the 41 common deregulated miRNAs, miR-125b-5p, let-7c-5p and miR-204-5p were found to be significantly downregulated in OSCC and abundantly expressed in normal tissue samples. These three miRNAs also showed significant associations with patient survival by acting as potent tumor suppressors in oral cancer. The target genes of these miRNAs were found to regulate important cellular pathways in cancer such as, MAPK signaling, JAK-STAT signaling, p53 signaling, and ECM receptor signaling. The common oncogenic targets of these 3 miRNA-pairs that were upregulated in oral cancer were identified and validated using luciferase assays. Transfection of the individual recombinant miRNA-plasmids or miRNA mimics downregulated the 3’-UTR target genes. Overexpression of the miRNA-pairs with one-half of the dose of the recombinant miRNA-plasmids or miRNA mimics resulted in a greater reduction in luciferase activity, suggesting a probable synergistic interaction between the miRNA-pairs and some of the UTRs. We observed that the distance between the MREs on the 3’-UTR of their common target genes has an important role in the synergistic downregulation of gene expression. Generally, a distance ranging from 200bp to 800bp showed synergistic effect with co-expression of the miRNA-pairs. However, some of the target UTRs with distances between their MREs greater than or less than the distance range also exhibited synergistic effects, suggesting the involvement of other factors, such as the RNA secondary structure, Gibbs free energy of binding and conservation of the MREs in miRNA-mediated synergism. To further validate the specific role of the distance between MREs on synergism, insertion/deletion constructs with varying distances between the MREs were used. Since, the insertion/deletion plasmids were constructed using the same 3’-UTR recombinant plasmids, the effect of other regulatory factors on the downregulation of gene expression were eliminated, and the results truly reflected the effect of the distance between MREs on miRNA-mediated synergism. Co-transfection of the insertion/deletion plasmids along with a half dose of each of the miRNAs also revealed that an optimal distance between 200bp and 800bp showed synergistic effect. Moreover, deletion of either of the 6bp seed sequences of let-7c-5p or miR-125b-5p in the MRE of the 3’-UTRs of HK2 and BCAT1 constructs resulted in a loss of synergistic activity strengthening our hypothesis of miRNA-mediated co-operative binding and synergistic effects.

To study the actual effect of miRNA-mediated synergism in cancer and study its effect on cellular toxicity and off-target effects, ORL48 oral cancer cells were transfected with either 20pmol of the individual miRNA mimics or 10pmol of the combination of miRNA mimic-pairs. A reduction in the expression of common and unique target genes was observed with the over-expression of the miRNA mimics individually. Moreover, co-expressing a half dose of the miRNA-pairs significantly and synergistically reduced the expression of common target genes, while the unique target genes did not show any significant reduction in their expression. A reduction in HK2 and BCAT1 protein expression was observed with overexpression of the miRNAs individually or in combination compared to that in the mock control, but, a greater fold reduction was observed with co-transfection of the miRNA-pairs indicating a synergistic effect. On the contrary, both HK2 and BCAT1 did not show any synergistic reduction in their protein expression upon co-expression of the miRNAs in the normal keratinocyte cells. HK2 and BCAT1 are oncogenes and are reported to be highly expressed in oral cancer, while, these two genes are expressed at low levels under normal condition and are required for normal cell growth and metabolism. Therefore, co-expression of miRNAs results in a synergistic reduction in their expression in cancer cells owing to their greater abundance in cancer but does not alter their expression in normal cell-lines.

HK2 and BCAT1 were identified as crucial targets of let-7c-5p and miR-125b-5p that showed synergistic down-regulation of their gene and protein expression with the co-expression of the miRNAs. To further study the effect of miRNA-mediated synergism on the function and metabolism of oral cancer cells, the global proteomic and metabolomic profiles of oral cancer cells were analyzed. Transfection of individual miRNA mimics or half doses of miRNA mimic-pairs in the oral cancer cells significantly reduced the expression of several proteins. Among the 6 common proteins targeted by let-7c-5p and miR-125b-5p, HK2, BCAT1, and STAT3 showed significant synergistic downregulation of their protein expression after co-transfection with a half dose of the miRNA-pairs and were also found to be upregulated in oral cancer. The remaining 3 proteins, AIFM1, MAP3K9, and G3BP1 did not exhibit any significant synergistic downregulation of their expression. The distances between the MREs of the miRNA-pairs in the HK2, BCAT1 and STAT3 3’-UTRs were found to be 401bp, 594bp, and 1448bp respectively, while, the distances between the MREs of AIFM1, MAP3K9, and G3BP1 were found to be -3bp, 1178bp, and 2379bp, respectively. Since, the HK2 and BCAT1 proteins showed significant synergistic effect while, AIFM1, MAP3K9, and G3BP1 did not show significant synergistic effect, these findings further validate our hypothesis that an optimal distance between 200bp and 800bp on the MREs of the target gene shows synergistic effect. In contrast, STAT3 was significantly synergistically downregulated despite having a distance of 1448bp between the MREs of the miRNA-pair, suggesting that other contributing factors play important roles in the regulation of miRNA-mediated synergism and is an open area of research. Moreover, the effect of the other abundantly expressed miRNAs that may have MREs at the 3’-UTR of the target gene, and the abundance of the target gene in a particular cell or tissue might also play important roles in the regulation of synergism.

The unique target proteins of let-7c-5p and miR-125b-5p upregulated in oral cancer, namely, IGF2BP3, GALNT2, CALU, HMGB3, and PSMB9, did not show any significant down-regulation in their protein expression with co-expression of the miRNAs, indicating target specificity and a reduction in off-target effects (**Figure 4e**). Among the remaining proteins that were predicted to be uniquely targeted by the respective miRNAs, many did not show any target specific deregulation. Although, some of the proteins targeted by either let-7c-5p or miR-125b-5p were significantly downregulated in the respective miRNA-transfected cells, all these proteins were also significantly downregulated in cells transfected with the other miRNA, suggesting miRNA independent downstream effect for these proteins.

HK2 catalyzes the first rate-determining step of glycolysis, resulting in the conversion of glucose to glucose-6-phosphate[34]. The metabolomic profile showed a reduction in glucose-phosphate as well as an increase in glucose and the glucose-to-glucose-phosphate ratio in cancer cells transfected with individual miRNAs or miRNA mimic-pairs. A greater abundance of glucose and a greater reduction in glucose-phosphate were observed with co-transfection of the miRNA-pair than with the individual mimic transfected cells, indicating the synergistic effects of the effects of the miRNA mimic-pairs. Moreover, overall decreases in glycolysis, the TCA cycle, the pentose phosphate pathway and oxidative phosphorylation were observed in cells co-transfected with a half dose of the miRNA mimic-pair compared to the individual mimic transfected cells, further validating the effectiveness of this approach. A synergistic reduction in BCAT1 was also observed in the proteomic data. BCAT1 catalyzes the conversion of BCAAs to α-keto acids [35]. A reduction in BCAT1 resulted in a decrease in BCAA degradation and an increase in the synthesis of valine, leucine, and isoleucine. A reduction in BCAT1 was also evident from the metabolomic data where an increase in the BCAA isoleucine level was observed with the co-expression of miR-125b-5p and let-7c-5p in oral cancer cells. Moreover, pathway enrichment analysis of the significant genes revealed a reduction in HIF-signaling, IL-17 signaling and other important oncogenic pathways activated in cancer in the co-transfected cells compared to the individual miRNA mimic transfected cells. Taken together, these findings indicate that the co-expression of miRNAs might specifically target oncogenic proteins and effectively alter the cancer phenotype but reduce off-target effects, thus, reducing cellular toxicity.

Finally, to understand the effect of miRNA synergy on the cancer phenotype, oral cancer cells were transfected with either the individual miRNAs or a half dose of the combination of each miRNA. A reduction in cell proliferation and migration was observed following the transfection of the miRNAs alone or in combination. A greater fold reduction in cell proliferation and migration rate was observed with co-transfection of the miRNA-pairs. However, co-transfection of the miRNA-pair did not alter the phenotype of the normal cells. Hence, co-expression of miRNAs with reduced doses effectively downregulates the common oncogenic targets in cancer cells, effectively altering the cancer phenotype but does not affect the normal cells, thus reducing cellular toxicity. A reduction in the diameter of cancer spheroids was also observed with co-transfection of the miRNAs in the oral cancer cells compared to the mock controls as well as the individual mimic transfected cells. Since, 3D-spheroids are physiologically more relevant than 2-D formats in terms of recapitulating the features of native tumor microenvironments, a reduction in spheroid formation with miRNA co-transfection in cancer cells is suggestive of the potential of this approach in therapeutics.

The present study investigated the mechanism of miRNA-mediated synergism, and applied this approach in oral cancer. Our study suggested that miRNA synergism-based therapeutics could be promising tools for eradicating the drawbacks of conventional miRNA therapeutics. However, the contributions of other factors, such as target gene abundance, 3’-UTR RNA secondary structure, selection of miRNA pairs and the abundance of other miRNAs with their binding sites in the target gene require further investigation followed by validation in animal models.

## ACKNOWLEDGEMENTS

We would like to thank all the members of the Human Genetics Unit, ISI and all the doctors who provided us with the samples for their co-operation. We would also like to extend our gratitude to the patients who participated in the study. We would like to thank Prof. Sok Ching Cheong (Department of Oro-Maxillofacial Surgery and Medical Sciences, University of Malaya) for providing us with the ORL-48 cell-line. SM is also thankful to CSIR for funding her research fellowship.

## AUTHOR CONTRIBUTIONS

S.M. and R.C designed the research; S.M., U.B., S.S., S.M., D.P., and P.B. performed the research; S.M., D.P., S.M., P.B., B.D. and R.C analyzed the data; and S.M. and R.C wrote the paper. All the authors edited and approved the manuscript.

## Author Declarations

All the patient tissue samples used for this study have been collected and used *vide ISI Ethics Clearance and Review Committee for Protection of Research Risks to Humans (*ISI-IEC/2022/07/04 dated July 28, 2022).

## DATA AVAILABILITY

Small RNA-Sequencing data of 4 OSCC and 4 adjacent normal tissue samples are available in the ***Gene Expression Omnibus database*** (https://www.ncbi.nlm.nih.gov/geo): ***Accession Number:*** GSE270895

The publicly available ***“The Cancer Genome Atlas (TCGA)” small RNA-sequencing data of HNSCC*** was downloaded from ***TCGA-GDC Data Portal (NIH)*** (https://portal.gdc.cancer.gov). The manifest files and the clinical data files for each patient was downloaded using GDC Data Transfer tool (https://github.com/NCI-GDC/gdc-client).

The publicly available “The Cancer Genome Atlas (TCGA)” RNA-sequencing data of HNSCC was downloaded from TCGA-GDC Data Portal (NIH) (https://portal.gdc.cancer.gov).

The publicly available ***“Gene expression profile of OSCC, oral dysplasia, and normal oral tissue”*** was downloaded from the ***EMBL-EBI Expression Atlas*** (https://www.ebi.ac.uk).

The target genes of the miRNAs were predicted bioinformatically using ***Target Scan Human 7.1 (Agarwal et al., 2015)*** software.

