## Supplementary Tables for "miRNA synergism impaired metabolic programming of oral cancer cells by selectively enhancing target gene specificity"

**Table S1:** Demographic information of oral squamous cell carcinoma patients included in the discovery set

| Sample ID | Age (in years) | Sex | Histopathological diagnosis |
|---|---|---|---|
| DC105 | 66 | Male | Well differentiated Carcinoma |
| DC106 | 35 | Male | Well differentiated Carcinoma |
| DC125 | 35 | Male | Well differentiated Carcinoma |
| DC16_11 | 55 | Male | Well differentiated Carcinoma |

**Table S2:** Demographic information of oral squamous cell carcinoma patients included in the validation set

| Sample ID | Age (in years) | Sex | Histopathological diagnosis |
|---|---|---|---|
| DC200 | 55 | Male | Well differentiated carcinoma |
| DC205 | 73 | Male | Well differentiated carcinoma |
| DC206 | 60 | Male | Well differentiated carcinoma |
| DC112 | 60 | Male | Well differentiated carcinoma |
| DC220 | 60 | Male | Well differentiated carcinoma |
| DC105 | 66 | Male | Well differentiated Carcinoma |
| DC106 | 35 | Male | Well differentiated Carcinoma |
| DC16-32 | | | Well differentiated carcinoma |
| DC211 | 60 | Male | Well differentiated carcinoma |
| DC212 | 32 | Male | Well differentiated carcinoma |
| DC103 | 65 | Female | Well differentiated carcinoma |
| DC92 | 41 | Male | Well differentiated carcinoma |
| DC94 | 48 | Female | Well differentiated carcinoma |
| DC104 | 70 | Female | Well differentiated carcinoma |
| DC18 | 45 | Male | Well differentiated carcinoma |
| DC34 | 36 | Male | Well differentiated carcinoma |
| DC97 | 60 | Male | Well differentiated carcinoma |
| DC6 | 52 | Male | Well differentiated carcinoma |
| DC12 | 70 | Male | Well differentiated carcinoma |
| DC32 | 54 | Female | Well differentiated carcinoma |
| DC39 | 67 | Male | Well differentiated carcinoma |
| DC45 | 55 | Male | Well differentiated carcinoma |
| DC62 | 48 | Male | Well differentiated carcinoma |
| DC52 | 56 | Female | Well differentiated carcinoma |
| DC69 | 44 | Male | Well differentiated carcinoma |
| DC70 | 80 | Female | Well differentiated carcinoma |
| DC57 | 44 | Male | Well differentiated carcinoma |
| DC116 | 63 | Male | Well differentiated carcinoma |
| DC120 | 45 | Male | Well differentiated carcinoma |
| DC58 | 70 | Female | Well differentiated carcinoma |
| DC59 | 40 | Male | Well differentiated carcinoma |
| DC73 | 42 | Female | Well differentiated carcinoma |
| DC77 | 45 | Male | Well differentiated carcinoma |

**Table S3:** Illumina small RNA sequencing data alignment summary

| Sample ID | Total Reads | Reads remaining after quality trimming | Total Mapped | | | |
|---|---|---|---|---|---|---|
| | | | Uniquely mapped | Multi-mapped | No mapping found | QC failure |
| mi3_OSCC | 18422652 | 12278015 | 4917849 (40.1%) | 3969149 (32.3%) | 2149903 (17.5%) | 1241114 (10.1%) |
| mi4_OSCC | 23647492 | 15851845 | 8494796 (53.6%) | 3297560 (20.8%) | 2763912 (17.4%) | 1295577 (8.2%) |
| mi5_OSCC | 17284107 | 12554481 | 5283385 (42.1%) | 4261083 (33.9%) | 2104552 (16.8%) | 905461 (7.2%) |
| mi6_OSCC | 19465777 | 12225348 | 5521548 (45.2%) | 3558805 (29.1%) | 1943363 (15.9%) | 1201632 (9.8%) |
| mi9_OSCC | 21489427 | 16562708 | 6852803 (41.4%) | 5685305 (34.3%) | 2897084 (17.5%) | 1127516 (6.8%) |
| mi10_OSCC | 19382352 | 14734743 | 7450755 (50.6%) | 4581601 (31.1%) | 1809804 (12.3%) | 892583 (6.1%) |
| mi7_OSCC | 40118921 | 39505134 | 12549687 (31.8%) | 15022413 (38.0%) | 11414720 (28.9%) | 518314 (1.3%) |
| mi8_OSCC | 29514970 | 20617347 | 10512001 (51.0%) | 5137009 (24.9%) | 3120272 (15.1%) | 1848065 (8.9%) |

**Table S4:** Small RNA sequencing data. Significantly differentially regulated miRNAs between OSCC and adjacent normal tissue

| miRNA | logFC | logCPM | LR | PValue | FDR |
|---|---|---|---|---|---|
| hsa-miR-31-3p | 6.567127 | 6.651854 | 35.33019 | 2.78E-09 | 9.68E-07 |
| hsa-miR-31-5p | 6.172747 | 14.61467 | 26.90361 | 2.14E-07 | 2.98E-05 |
| hsa-miR-196a-5p | 5.869728 | 5.823671 | 15.62485 | 7.72E-05 | 0.003162 |
| hsa-miR-767-5p | 4.426442 | 3.118252 | 9.23077 | 0.00238 | 0.026715 |
| hsa-miR-2355-3p | 3.500968 | 2.186815 | 8.429341 | 0.003692 | 0.036193 |
| hsa-miR-937-3p | 3.059674 | 2.569953 | 14.04073 | 0.000179 | 0.004789 |
| hsa-miR-503-5p | 2.986415 | 4.862287 | 14.70442 | 0.000126 | 0.003775 |
| hsa-miR-21-3p | 2.841903 | 8.178714 | 11.5567 | 0.000675 | 0.011746 |
| hsa-miR-135b-3p | 2.834727 | 3.787405 | 14.8683 | 0.000115 | 0.003775 |
| hsa-miR-2355-5p | 2.80662 | 2.87078 | 10.05649 | 0.001518 | 0.019936 |
| hsa-miR-21-5p | 2.748881 | 18.5711 | 15.9618 | 6.46E-05 | 0.003162 |
| hsa-miR-4664-3p | 2.729575 | 2.063926 | 13.58213 | 0.000228 | 0.00548 |
| hsa-miR-944 | 2.535955 | 8.154862 | 16.22985 | 5.61E-05 | 0.003004 |
| hsa-miR-196b-5p | 2.52907 | 5.749878 | 10.29374 | 0.001335 | 0.018581 |
| hsa-miR-301b | 2.452084 | 3.795032 | 9.830603 | 0.001716 | 0.021719 |
| hsa-miR-431-5p | 2.352775 | 4.300434 | 10.10129 | 0.001482 | 0.019894 |
| hsa-miR-708-3p | 2.348976 | 6.565726 | 13.68644 | 0.000216 | 0.005369 |
| hsa-miR-7705 | 2.152297 | 2.791516 | 8.56161 | 0.003433 | 0.034136 |
| hsa-miR-223-3p | 2.136945 | 8.759013 | 10.61938 | 0.001119 | 0.016572 |
| hsa-miR-455-3p | 2.071333 | 8.118715 | 9.029284 | 0.002657 | 0.028449 |
| hsa-miR-181b-5p | 1.894568 | 9.34658 | 9.360238 | 0.002217 | 0.025301 |
| hsa-miR-7-5p | 1.89288 | 5.768515 | 8.859795 | 0.002915 | 0.029838 |
| hsa-miR-181b-3p | 1.864824 | 4.048691 | 8.91272 | 0.002832 | 0.029418 |
| hsa-miR-101-3p | -2.01701 | 7.725086 | 7.784044 | 0.005271 | 0.047644 |
| hsa-miR-653-5p | -2.04502 | 3.202769 | 9.674915 | 0.001868 | 0.022888 |
| hsa-miR-139-3p | -2.04897 | 5.014768 | 10.33307 | 0.001307 | 0.01856 |

| | | | | | |
|---|---|---|---|---|---|
| hsa-miR-30a-3p | -2.14724 | 9.785556 | 11.19643 | 0.00082 | 0.012964 |
| hsa-miR-99a-3p | -2.17345 | 6.10867 | 8.744352 | 0.003106 | 0.031326 |
| hsa-miR-144-5p | -2.19543 | 9.140271 | 8.057351 | 0.004532 | 0.042625 |
| hsa-miR-30c-2-3p | -2.31177 | 6.274626 | 13.96306 | 0.000186 | 0.004806 |
| hsa-miR-30a-5p | -2.40068 | 14.44331 | 11.86789 | 0.000571 | 0.01046 |
| hsa-miR-296-5p | -2.51845 | 2.258159 | 7.720299 | 0.00546 | 0.048723 |
| hsa-miR-1247-5p | -2.57481 | 3.311933 | 11.66012 | 0.000639 | 0.011396 |
| hsa-miR-20b-5p | -2.69844 | 6.129677 | 10.72465 | 0.001057 | 0.015996 |
| hsa-miR-504-5p | -2.70211 | 5.126025 | 12.09229 | 0.000506 | 0.009524 |
| hsa-miR-125b-2-3p | -2.70866 | 7.649878 | 18.8207 | 1.44E-05 | 0.000999 |
| hsa-miR-151b | -2.77569 | 2.714069 | 14.74096 | 0.000123 | 0.003775 |
| hsa-miR-144-3p | -2.81056 | 8.97755 | 9.168971 | 0.002462 | 0.027046 |
| hsa-miR-100-5p | -2.82929 | 12.45857 | 11.2528 | 0.000795 | 0.012868 |
| hsa-miR-363-3p | -2.83728 | 8.118615 | 9.667195 | 0.001876 | 0.022888 |
| hsa-miR-451b | -2.99459 | 2.425392 | 7.673883 | 0.005603 | 0.049359 |
| hsa-let-7c-5p | -3.02081 | 13.10085 | 19.66695 | 9.22E-06 | 0.000713 |
| hsa-miR-335-5p | -3.12622 | 8.211828 | 12.67527 | 0.000371 | 0.007368 |
| hsa-miR-328-3p | -3.24876 | 6.592322 | 14.56224 | 0.000136 | 0.003775 |
| hsa-miR-508-3p | -3.64798 | 4.31417 | 21.48128 | 3.57E-06 | 0.000355 |
| hsa-miR-375 | -3.69468 | 11.05832 | 15.23968 | 9.47E-05 | 0.003661 |
| hsa-let-7c-3p | -3.78892 | 2.680513 | 20.71811 | 5.32E-06 | 0.000463 |
| hsa-miR-513c-5p | -4.12262 | 2.295225 | 13.16643 | 0.000285 | 0.006421 |
| hsa-miR-99a-5p | -4.18274 | 13.81088 | 14.59417 | 0.000133 | 0.003775 |
| hsa-miR-125b-5p | -4.29452 | 6.449582 | 10.80011 | 0.001015 | 0.015698 |
| hsa-miR-6510-3p | -4.35208 | 3.349461 | 28.5816 | 8.98E-08 | 1.56E-05 |
| hsa-miR-204-5p | -4.76105 | 6.599856 | 44.88812 | 2.09E-11 | 1.45E-08 |
| hsa-miR-509-3-5p | -4.98589 | 4.310132 | 15.77248 | 7.14E-05 | 0.003162 |
| hsa-miR-891a-5p | -5.04008 | 3.110444 | 18.1455 | 2.05E-05 | 0.001295 |
| hsa-miR-6744-5p | -7.14723 | 2.505604 | 14.57619 | 0.000135 | 0.003775 |
| hsa-miR-383-5p | -7.77652 | 2.104058 | 26.24329 | 3.01E-07 | 3.49E-05 |
| hsa-miR-211-5p | -7.78195 | 5.141804 | 32.72314 | 1.06E-08 | 2.47E-06 |

**Table S5:** Expression of the Clinically Significant miRNAs in TCGA-HNSC and small RNA-sequencing data

| **MiRNAs** | **Log(FC) [log(CPM) in OSCC/HNSCC]** | | | | |
|---|---|---|---|---|---|
| | **Indian OSCC** | **TCGA Stage1** | **TCGA Stage2** | **TCGA Stage3** | **TCGA Stage4a** |
| miR-204-5p | -4.7611 | -2.041 | -3.2287 | -2.6439 | -2.0567 |
| | [6.5998] | [3.4659] | [2.9647] | [2.9102] | [2.7220] |
| miR-125b-5p | -4.2945 | -1.0067 | -1.0496 | -1.0623 | -1.1415 |
| | [6.4496] | [8.3501] | [8.2754] | [8.1339] | [7.9727] |
| Let-7c-5p | -3.0208 | -1.8913 | -2.4046 | -1.8969 | -2.1819 |
| | [13.1008] | [12.0526] | [11.6846] | [11.6522] | [11.2181] |
| miR-99a-5p | -4.1827 | -2.2073 | -2.6201 | -2.0158 | -2.396 |
| | [13.8109] | [9.9272] | [9.5526] | [9.5378] | [9.0212] |
| miR-99a-3p | -2.1734 | -2.2073 | -2.6201 | -2.0158 | -2.396 |
| | [6.1087] | [9.9272] | [9.5526] | [9.5378] | [9.0212] |
| miR-100-5p | -2.8293 | -1.5392 | -1.5454 | -1.8173 | -1.7226 |
| | [12.4586] | [12.3239] | [12.1564] | [11.9033] | [11.7072] |
| miR-125b-2-3p | -2.7087 | -1.1016 | -1.1454 | -1.1427 | -1.2256 |
| | [7.6498] | [8.5298] | [8.4371] | [8.2985] | [8.1197] |
| miR-2355-3p | 2.8066 | 1.3377 | 1.4351 | 1.6057 | 1.5097 |
| | [2.8708] | [5.8412] | [6.3569] | [6.4481] | [6.6553] |

**Table S6:** KEGG Pathway Analysis for Union of Genes Targeted by Let-7c-5p, miR-125b-5p and miR-204-5p

| **Pathway** | **P-value** |
|---|---|
| ECM-receptor interaction | 7.36E-14 |
| PI3K-Akt signaling pathway | 3.82E-06 |
| Glycosaminoglycan biosynthesis - chondroitin sulfate | 9.91E-06 |
| Amoebiasis | 3.83E-05 |
| MAPK signaling pathway | 4.27E-04 |
| Valine, leucine, and isoleucine biosynthesis | 7.72E-04 |
| Protein digestion and absorption | 1.19E-03 |
| Focal adhesion | 1.46E-03 |
| Neurotrophin signaling pathway | 0.002242 |
| Transcriptional mis regulation in cancer | 0.014236 |
| Ubiquitin mediated proteolysis | 0.017418 |
| Cytokine-cytokine receptor interaction | 0.022489 |
| Dilated cardiomyopathy | 0.025415 |
| HTLV-I infection | 0.029236 |
| Jak-STAT signaling pathway | 0.031853 |

**Table S7:** Distance between MRE *versus* synergism ($E_{syn}$) with co-transfection of recombinant pmirGLO plasmid and recombinant pRNA-U6 plasmids or miRNA mimics

| Gene | Distance | Synergism on mimic transfection | Synergism on pRNA-U6 plasmid transfection |
|---|---|---|---|
| NKIRAS2 | 488 | 1.5 | 4.8 |
| HK2 | 401 | 1.1 | 7.2 |
| CCDC71L | 180 | 1.3 | 1.2 |
| NIPAL4 | 475 | 1.2 | 1.1 |
| BCAT1 | 594 | 1.2 | 1.2 |
| USP38 | 534 | 1.2 | 2.7 |
| BCL2L2 | 109 | 1.5 | 1.8 |
| PTPRD | 816 | 1.4 | 1.2 |
| HAS2 | 622 | 1.4 | 2.1 |
| HMGA2 | 584 | 1.9 | 3.3 |
| MEX3A | 137 | 0.2 | 1.4 |
| ANGPTL2 | 45 | 0.3 | 0.1 |
| LRRC10B | 630 | 2.3 | 2 |
| BTG2 | 410 | 2.3 | 2.3 |
| PTAR1 | 307 | 4.2 | 4.6 |
| C15ORF39 | 465 | 2.3 | 4.9 |
| ELOVL4 | 863 | 0 | 1.5 |
| CLDN12 | 941 | 0.4 | 0.2 |
| SEMA4C | 11 | 1.6 | 1.1 |
| ARID3B | 247 | 3.59 | 6.1 |
| ACER2 | 676 | 2.8 | 2.2 |
| TET3 | 612 | 1.84 | 2.1 |
| MIB1 | 377 | 2.32 | 1.5 |
| ERCC6 | 180 | 1.85 | 2.8 |
| ACVR2B | 423 | 1.98 | 1.9 |
| LBH | 750 | NA | NA |
| MTUS1 | 708 | NA | NA |
| SOX11 | 1318 | 0 | 0 |

**Table S8:** Distance between MRE *versus* synergism ($E_{syn}$) with co-transfection of insertion/deletion constructs and recombinant pRNA-U6 plasmids or miRNA mimics

| Gene | Distance | Synergism on mimic transfection | Synergism on pRNAU6 plasmid transfection |
|---|---|---|---|
| HK2_DEL1 | 101 | 0.6 | 0.5 |
| BCAT1_DEL1 | 145 | 0.9 | 0.5 |
| CCDC71L | 180 | 0.6 | 1.1 |
| HK2_DEL2 | 301 | 1.7 | 2.4 |
| HK2 | 401 | 4.8 | 1.3 |
| CCDC71L_IN1 | 450 | 1.8 | 1.6 |
| BCAT1 | 594 | 1.1 | 1.2 |
| HK2_IN1 | 640 | 1.7 | 1.5 |
| CCDC71L_IN2 | 700 | 1.3 | 1.1 |
| HK2_IN2 | 890 | 0.9 | 0.9 |
| CCDC71L_IN3 | 900 | 0.9 | 0.6 |
| HK2_IN3 | 1090 | 0.8 | 0.7 |

**Table S9:** Proteomics Data_List of Common and Unique Target Genes of Let-7c-5p and miR-125b-5p

| Accessions | FC_Let7c | FC_miR125b | FC_Both | p-value_7c | p-value_125b | p-value_Both |
|---|---|---|---|---|---|---|
| **Common targets of Let-7c and miR-125b** | | | | | | |
| HK2 | 0.887932 | 0.801667 | 0 | 0.3754227 | 0.0487138 | 0 |
| AIFM1 | 0.947837 | 1.242715 | 1.726685 | 0.6859853 | 0.2513872 | 0.001821051 |
| STAT3 | 0.734018 | 0.803173 | 0.44123 | 0.0053655 | 0.0759982 | 1.85159E-06 |
| BCAT1 | 0.774354 | 0.809549 | 0.620048 | 0.0405579 | 0.0744876 | 0.001829932 |
| MAP3K9 | 1.056721 | 1.021848 | 0.849066 | 0.5482223 | 0.7833046 | 0.082489145 |
| G3BP1 | 0.76231 | 0.866484 | 0.878618 | 0.0016558 | 0.0864516 | 0.076154495 |
| **Unique targets of Let-7c** | | | | | | |
| UFM1 | 1.391584 | 1.25515 | 1.832786 | 0.0289085 | 0.1542529 | 0.000202637 |
| IGF2BP2 | 0.352067 | 0.506462 | 0.149492 | 0.000673 | 0.0015373 | 0.000141512 |
| NRAS | 0.899903 | 0.864958 | 0.636627 | 0.2831417 | 0.0951891 | 3.24775E-05 |
| HMGA1 | 0.994346 | 1.088625 | 1.406527 | 0.9527848 | 0.4737427 | 0.004604933 |
| DPP3 | 0.865087 | 0.914487 | 0.675556 | 0.2517889 | 0.482156 | 0.003684264 |
| IGF2BP3 | 0.671853 | 0.615335 | 0.611522 | 0.0034993 | 0.0014912 | 0.001549275 |
| BTF3L4 | 1.168632 | 1.594258 | 1.747756 | 0.1268895 | 0.022828 | 0.005855419 |
| SLC25A24 | 0.923847 | 1.070574 | 0.670941 | 0.5276767 | 0.5457532 | 0.001468952 |
| RTCA | 0.709891 | 0.748795 | 0.340186 | 0.0314909 | 0.0689379 | 6.04127E-07 |
| SYNCRIP | 1.053205 | 1.076122 | 1.178349 | 0.3824191 | 0.2159118 | 0.000419419 |
| CDV3 | 1.130429 | 1.160146 | 1.782748 | 0.2249173 | 0.3583772 | 1.60005E-05 |
| RCN1 | 1.000398 | 1.03058 | 1.57865 | 0.9968839 | 0.8211896 | 0.007569523 |
| GALNT2 | 0.994155 | 0.864023 | 0.632154 | 0.9673564 | 0.2319536 | 0.000316461 |
| PPP3CA | 0.987127 | 0.828891 | 0.802016 | 0.8654328 | 0.0565306 | 0.002906732 |
| CTPS1 | 0.868577 | 0.996865 | 0.679575 | 0.0332206 | 0.9551088 | 6.97883E-06 |
| LINGO1 | 1.846719 | 1.7054 | 2.564119 | 0.0045312 | 0.0116148 | 3.60199E-11 |
| CALU | 1.406469 | 1.438925 | 2.452462 | 0.0219913 | 0.0677192 | 0.000240299 |
| NUP155 | 0.890801 | 0.968794 | 0.657145 | 0.4111092 | 0.8072312 | 0.000548373 |
| HNRNPA1 | 1.331526 | 1.376101 | 1.73889 | 0.0229377 | 0.0123912 | 1.993E-08 |
| LRRC59 | 0.869185 | 0.887355 | 0.707415 | 0.021324 | 0.0141482 | 2.14366E-06 |
| NLN | 0.853843 | 0.889993 | 0.582647 | 0.1852955 | 0.3857963 | 1.00599E-05 |
| SUB1 | 1.211846 | 1.151448 | 1.910031 | 0.4077082 | 0.4156287 | 3.18622E-06 |
| XPOT | 1.072242 | 0.743262 | 0.670292 | 0.5397291 | 0.0096023 | 0.00046746 |
| NAP1L1 | 0.927159 | 1.019207 | 1.246221 | 0.4482108 | 0.8573873 | 0.035419512 |
| **Unique targets of miR-125b** | | | | | | |
| TMEM123 | 1.61669 | 1.4122 | 1.955068 | 0.0104582 | 0.0321024 | 0.000604529 |
| B4GALT1 | 0.729426 | 0.960225 | 0.460411 | 0.0678043 | 0.7827704 | 0.004166432 |
| TMED9 | 1.269827 | 1.038126 | 1.409378 | 0.2333158 | 0.7859854 | 0.028490543 |
| RHOT2 | 0.607838 | 0.686141 | 0.361754 | 4.548E-05 | 0.0110032 | 7.75698E-07 |
| NDRG3 | 0.74758 | 0.66437 | 0.46692 | 0.0088147 | 0.0067124 | 8.2688E-07 |
| YES1 | 0.943792 | 0.867669 | 0.696694 | 0.5831965 | 0.2141517 | 0.001137593 |
| TXNRD1 | 0.958521 | 0.926929 | 0.818778 | 0.4577417 | 0.1321337 | 0.008880789 |
| HMGB3 | 1.035246 | 1.038837 | 1.342357 | 0.6341077 | 0.5377137 | 0.001851643 |
| NCLN | 1.203903 | 1.33723 | 1.305479 | 0.1846294 | 0.0227148 | 0.026280508 |

| | | | | | | |
|---|---|---|---|---|---|---|
| YWHAG | 0.998196 | 1.025391 | 1.241259 | 0.9872617 | 0.848851 | 0.036333307 |
| PTPN1 | 1.102472 | 0.867697 | 1.348132 | 0.4922267 | 0.3601753 | 0.034015684 |
| CORO1C | 0.87797 | 0.786154 | 0.645756 | 0.0916722 | 0.0106195 | 6.92995E-05 |
| PSMB9 | 0.54607 | 0.678784 | 0.589182 | 0.0461932 | 0.1183223 | 0.064170483 |
| MYO1E | 0.469639 | 0.433981 | 0.168398 | 0.0068686 | 0.0008881 | 2.7731E-08 |
| SCARB2 | 1.101203 | 0.816804 | 1.534821 | 0.5368625 | 0.2339376 | 0.027438857 |
| HNRNPR | 1.067125 | 1.033088 | 1.12973 | 0.1950318 | 0.4618908 | 0.00474339 |
| UBE2L3 | 1.192213 | 1.341382 | 1.618301 | 0.0852749 | 0.0236299 | 2.09928E-07 |
| MRPL17 | 0.824203 | 0.895215 | 0.602208 | 0.1130974 | 0.289079 | 0.000179953 |
| ACOT13 | 0.662909 | 0.648427 | 0.472894 | 0.0019476 | 0.0011244 | 3.47713E-05 |

**Table S10:** Proteomics: KEGG Pathway Analysis of the Top Significantly Enriched Pathways

| **Pathway Enrichment with co-expression of let-7c-5p and miR-125b-5p** | | | |
|---|---|---|---|
| **pathways** | **enrichment** | **pvalue** | **count** |
| Biosynthesis of amino acids | 6.01098309 | 2.47E-11 | 10.6 |
| Fructose and mannose metabolism | 5.93970661 | 2.76E-05 | 4.6 |
| Pentose phosphate pathway | 5.880309544 | 7.51E-05 | 4.1 |
| Galactose metabolism | 5.058330791 | 0.00057 | 3.2 |
| Glycolysis / Gluconeogenesis | 4.973396132 | 2.98E-07 | 6.5 |
| Pyruvate metabolism | 4.587475531 | 0.00011 | 4.0 |
| Fatty acid metabolism | 4.470410765 | 3.31E-05 | 4.5 |
| Oxidative phosphorylation | 4.388290705 | 8.45E-11 | 10.1 |
| Glutathione metabolism | 4.126533014 | 0.00014 | 3.9 |
| Valine, leucine and isoleucine degradation | 4.083548295 | 0.00058 | 3.2 |
| Citrate cycle (TCA cycle) | 3.920206363 | 0.01234 | 1.9 |
| Alanine, aspartate and glutamate metabolism | 3.708303316 | 0.00859 | 2.1 |
| Central carbon metabolism in cancer | 3.640191623 | 0.00024 | 3.6 |
| DNA replication | 3.266838636 | 0.02631 | 1.6 |
| Apoptosis | 3.049049393 | 3.44E-05 | 4.5 |
| Glucagon signaling pathway | 2.930995411 | 0.0005 | 3.3 |
| HIF-1 signaling pathway | 2.337737739 | 0.01249 | 1.9 |
| IL-17 signaling pathway | 1.896874047 | 0.10334 | 1.0 |
| Pathways in cancer | 1.516306235 | 0.01709 | 1.8 |

| **Pathway Enrichment with expression of let-7c-5p** | | | |
|---|---|---|---|
| **pathways** | **enrichment** | **pvalue** | **count** |
| Pentose phosphate pathway | 8.671802187 | 2.79E-05 | 4.6 |
| Biosynthesis of amino acids | 7.804621969 | 2.96E-10 | 9.5 |
| Fructose and mannose metabolism | 6.898024467 | 0.0004 | 3.4 |
| Galactose metabolism | 6.294049975 | 0.00167 | 2.8 |
| Glycolysis / Gluconeogenesis | 5.824344753 | 1.02E-05 | 5.0 |
| Pyruvate metabolism | 5.535192886 | 0.00055 | 3.3 |
| Fatty acid metabolism | 5.134619716 | 0.00041 | 3.4 |
| Citrate cycle (TCA cycle) | 4.335901094 | 0.0435 | 1.4 |
| Central carbon metabolism in cancer | 4.181047483 | 0.00158 | 2.8 |
| Oxidative phosphorylation | 4.125577533 | 1.02E-05 | 5.0 |
| HIF-1 signaling pathway | 3.58010182 | 0.00082 | 3.1 |

| | | | |
|---|---|---|---|
| Apoptosis | 2.890600729 | 0.00463 | 2.3 |
| Valine, leucine and isoleucine degradation | 2.709938184 | 0.15092 | 0.8 |
| IL-17 signaling pathway | 2.447686101 | 0.07367 | 1.1 |
| Glucagon signaling pathway | 2.127428107 | 0.12719 | 0.9 |
| Pathways in cancer | 1.96342691 | 0.00139 | 2.9 |
| Alanine, aspartate and glutamate metabolism | 0 | 0 | 0.0 |
| DNA replication | 0 | 0 | 0.0 |
| Glutathione metabolism | 0 | 0 | 0.0 |

| **Pathway Enrichment with expression of miR-125b-5p** | | | |
|---|---|---|---|
| **pathways** | **enrichment** | **pvalue** | **count** |
| Biosynthesis of amino acids | 7.822156863 | 2.28E-12 | 11.6 |
| Fructose and mannose metabolism | 7.618983957 | 1.56E-05 | 4.8 |
| Pentose phosphate pathway | 6.518464052 | 0.00044 | 3.4 |
| Glycolysis / Gluconeogenesis | 6.254389816 | 1.44E-07 | 6.8 |
| Galactose metabolism | 5.407020873 | 0.00311 | 2.5 |
| Oxidative phosphorylation | 5.003511853 | 1.15E-09 | 8.9 |
| Fatty acid metabolism | 4.901100791 | 0.00021 | 3.7 |
| Central carbon metabolism in cancer | 4.78907563 | 5.59E-05 | 4.3 |
| Pyruvate metabolism | 4.755110555 | 0.00122 | 2.9 |
| Citrate cycle (TCA cycle) | 4.656045752 | 0.01475 | 1.8 |
| Apoptosis | 3.724836601 | 1.56E-05 | 4.8 |
| Valine, leucine and isoleucine degradation | 3.492034314 | 0.02353 | 1.6 |
| Glutathione metabolism | 3.430770554 | 0.01501 | 1.8 |
| Glucagon signaling pathway | 3.39412681 | 0.00069 | 3.2 |
| HIF-1 signaling pathway | 3.331849253 | 0.00077 | 3.1 |
| IL-17 signaling pathway | 3.304290533 | 0.00221 | 2.7 |
| Pathways in cancer | 1.897558269 | 0.00106 | 3.0 |
| Alanine, aspartate and glutamate metabolism | 0 | 0 | 0.0 |
| DNA replication | 0 | 0 | 0.0 |
