## Supplementary Figures for "miRNA synergism impaired metabolic programming of oral cancer cells by selectively enhancing target gene specificity"

### SUPPLEMENTARY FIGURES AND TABLES

#### SUPPLEMENTARY FIGURE LEGENDS

**Figure S1:** Validation of the common differentially regulated miRNAs between our small RNA sequencing data and TCGA-HNSCC small RNA sequencing data by qRT-PCR; p*≤0.05, ns>0.05.

**Figure S2:** *The functional role of miRNAs in oral cancer.* **a)** Expression of miRNAs in the stable cell-lines (ORL48_miR125b, ORL48_Let7c and ORL48_miR204) compared to the scramble control cell-line (ORL48_scrm) and the wild-type cell-line (ORL48_WT). **b)** WST1 cell proliferation assay. **c)** Scratch wound-healing cell migration assay **d)** Annexin-FITC-PI apoptosis assay. p*≤0.05, ns>0.05.

**Figure S3:** *Validation of the common targets of the miRNA-pairs.* **a)** Venn-diagram representing the number of common target genes between the miRNA-pairs that are up-regulated in oral cancer. **b)** Dose kinetic study with co-transfection of 10ng recombinant pmirGLO-3'-UTR plasmid and increasing doses of recombinant pRNA-U6 plasmids (0ng, 25ng, 50ng, 75ng, 100ng). **c)** Luciferase assay for validation of the common target genes of the miRNAs. d) Luciferase assay representing the synergistic down-regulation of the common target genes upon co-expression of half the dose (25ng) of recombinant pRNA-U6 plasmids. p*≤0.05, ns>0.05.

**Figure S4:** *Dose kinetic study.* **a)** Co-transfection of 10ng HK2_pmirGLO/BCAT1_pmirGLO plasmids and increasing doses of miRNA mimics (0pmol, 10pmol, 15pmol, 20pmol, 25pmol, 30pmol). **b)** Validation of optimal dose of miRNA mimics with co-transfection of NKIRAS2, NIPAL4, USP38, and CCDC71L 3'-UTR constructs and increasing doses of miRNA mimics (0pmol, 5pmol, 10pmol, 15pmol, 20pmol).

**Figure S5:** *Metabolomic assay.* **a-d)** Heatmap representing the enrichment of different metabolic pathways **e)** Relative abundance of metabolites in ORL-48 cells upon co-expression of half the dose (10pmol) of miRNA mimics. NC: Negative control mimic; S1: ORL-48 cells transfected with 20pmol NC mimic; S2: ORL-48 cells transfected with 20pmol let-7c-5p mimic; S3: ORL-48 cells transfected with 20pmol miR-125b-5p mimic; S4: ORL-48 cells transfected with 10pmol each of the mimic pair; p*≤0.05, ns>0.05.

### Figure S1

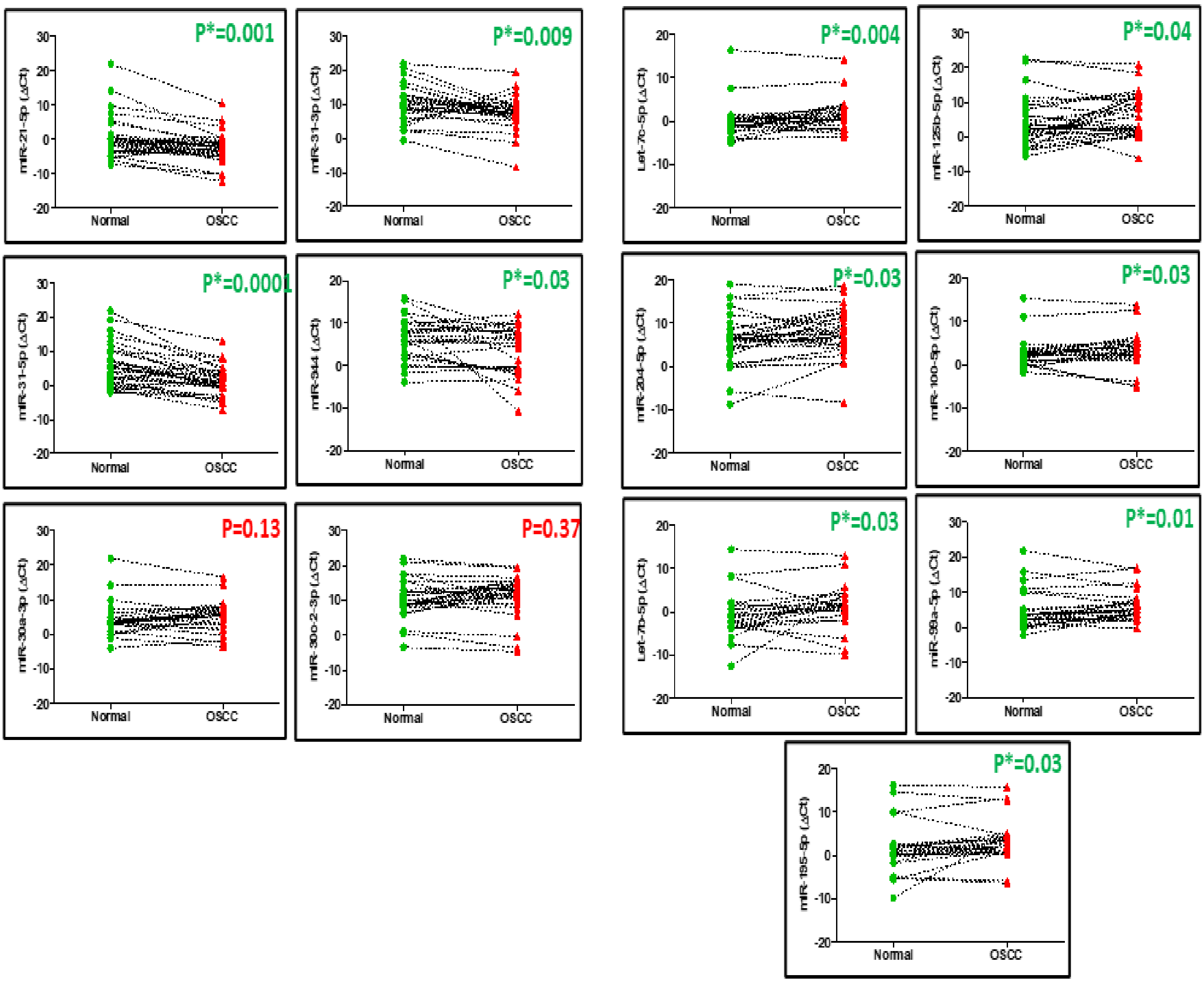

P*=0.001
mIR-21-5p (ΔCt)
Normal
OSCC
P*=0.009
mIR-31-3p (ΔCt)
Normal
OSCC
P*=0.004
Let-7c-5p (ΔCt)
Normal
OSCC
P*=0.04
mIR-125b-5p (ΔCt)
Normal
OSCC
P*=0.0001
mIR-31-5p (ΔCt)
Normal
OSCC
P*=0.03
mIR-944 (ΔCt)
Normal
OSCC
P*=0.03
mIR-204-5p (ΔCt)
Normal
OSCC
P*=0.03
mIR-100-5p (ΔCt)
Normal
OSCC
P=0.13
mIR-30a-3p (ΔCt)
Normal
OSCC
P=0.37
mIR-30c-2-3p (ΔCt)
Normal
OSCC
P*=0.03
Let-7b-5p (ΔCt)
Normal
OSCC
P*=0.01
mIR-99a-5p (ΔCt)
Normal
OSCC
P*=0.03
mIR-195-5p (ΔCt)
Normal
OSCC

### Figure S2

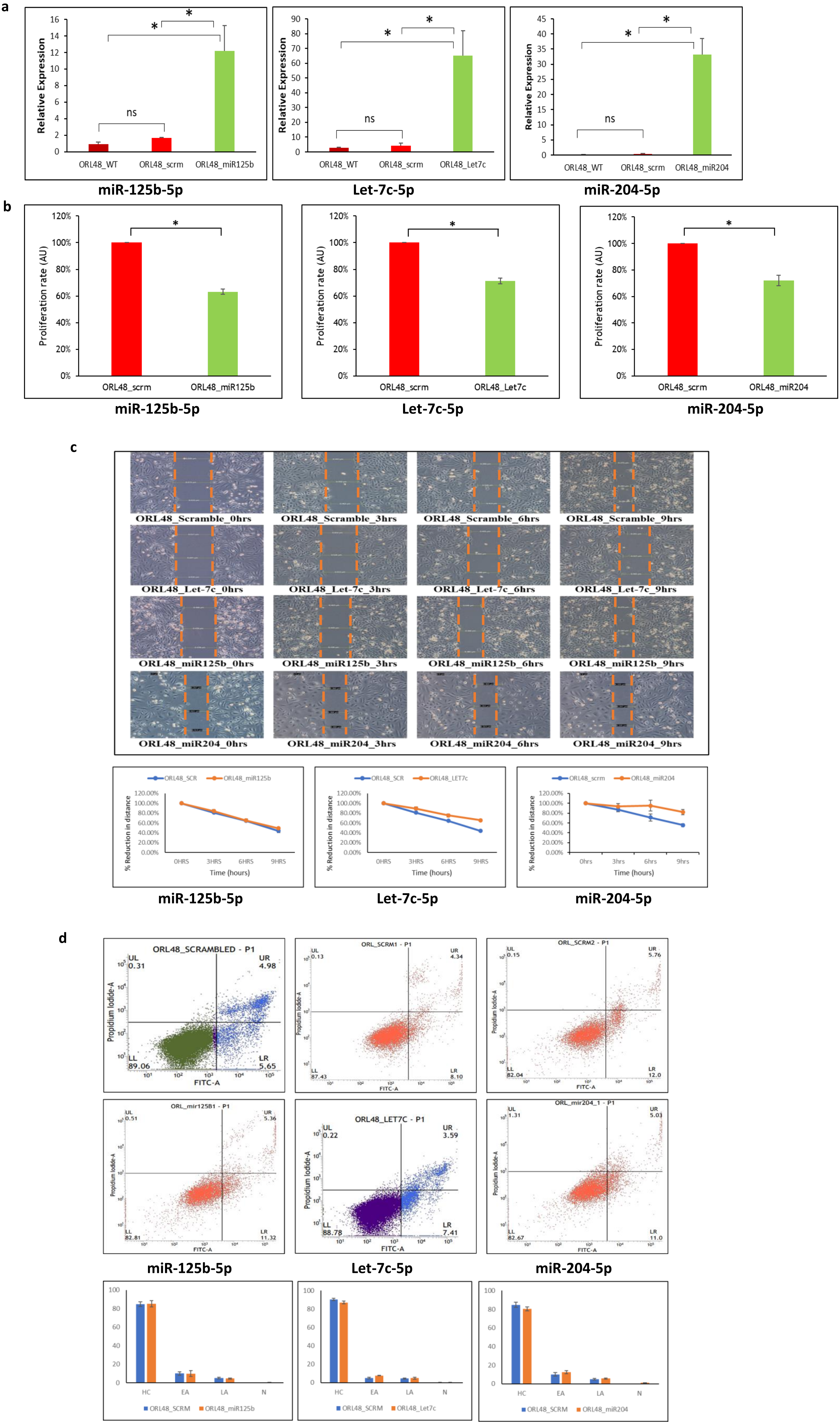


HC: Healthy Cells; EA: Early Apoptotic; LA: Late Apoptotic; N: Necrotic

### Figure S3

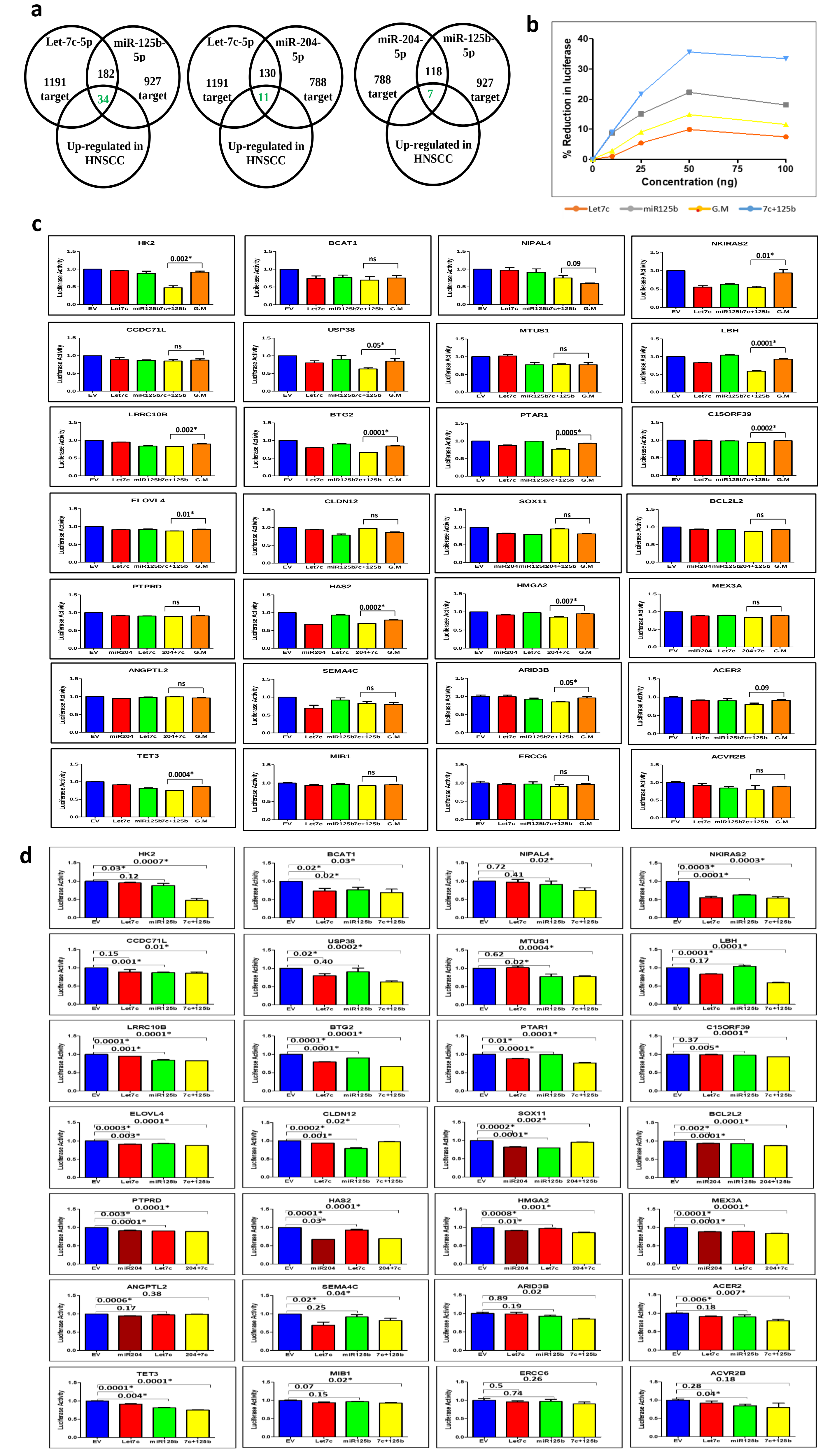
a
Let-7c-5p
miR-125b-5p
1191 target
182
927 target
34
Up-regulated in HNSCC
Let-7c-5p
miR-204-5p
1191 target
130
788 target
11
Up-regulated in HNSCC
miR-204-5p
miR-125b-5p
788 target
118
927 target
7
Up-regulated in HNSCC
b
% Reduction in luciferase
Concentration (ng)
Let7c
miR125b
G.M
7c+125b
c
HK2
BCAT1
NIPAL4
NKIRAS2
CCDC71L
USP38
MTUS1
LBH
LRRC10B
BTG2
PTAR1
C15ORF39
ELOVL4
CLDN12
SOX11
BCL2L2
PTPRD
HAS2
HMGA2
MEX3A
ANGPTL2
SEMA4C
ARID3B
ACER2
TET3
MIB1
ERCC6
ACVR2B
Luciferase Activity
d
HK2
BCAT1
NIPAL4
NKIRAS2
CCDC71L
USP38
MTUS1
LBH
LRRC10B
BTG2
PTAR1
C15ORF39
ELOVL4
CLDN12
SOX11
BCL2L2
PTPRD
HAS2
HMGA2
MEX3A
ANGPTL2
SEMA4C
ARID3B
ACER2
TET3
MIB1
ERCC6
ACVR2B
Luciferase Activity

### Figure S4

a

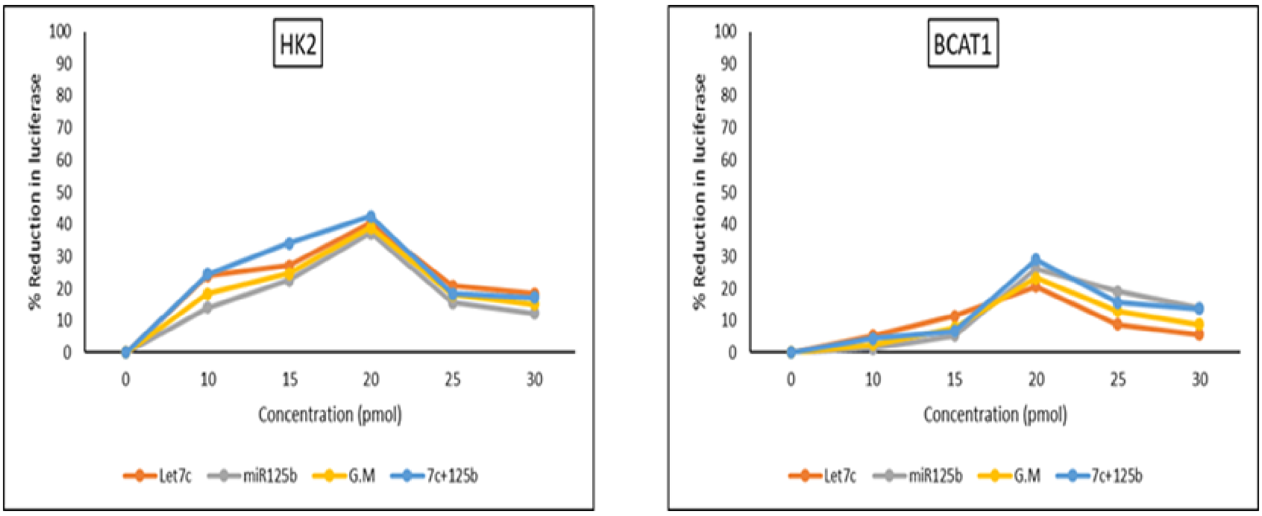
HK2
BCAT1
% Reduction in luciferase
100
90
80
70
60
50
40
30
20
10
0
0
10
15
20
25
30
Concentration (pmol)
Let7c
miR125b
G.M
7c+125b

b

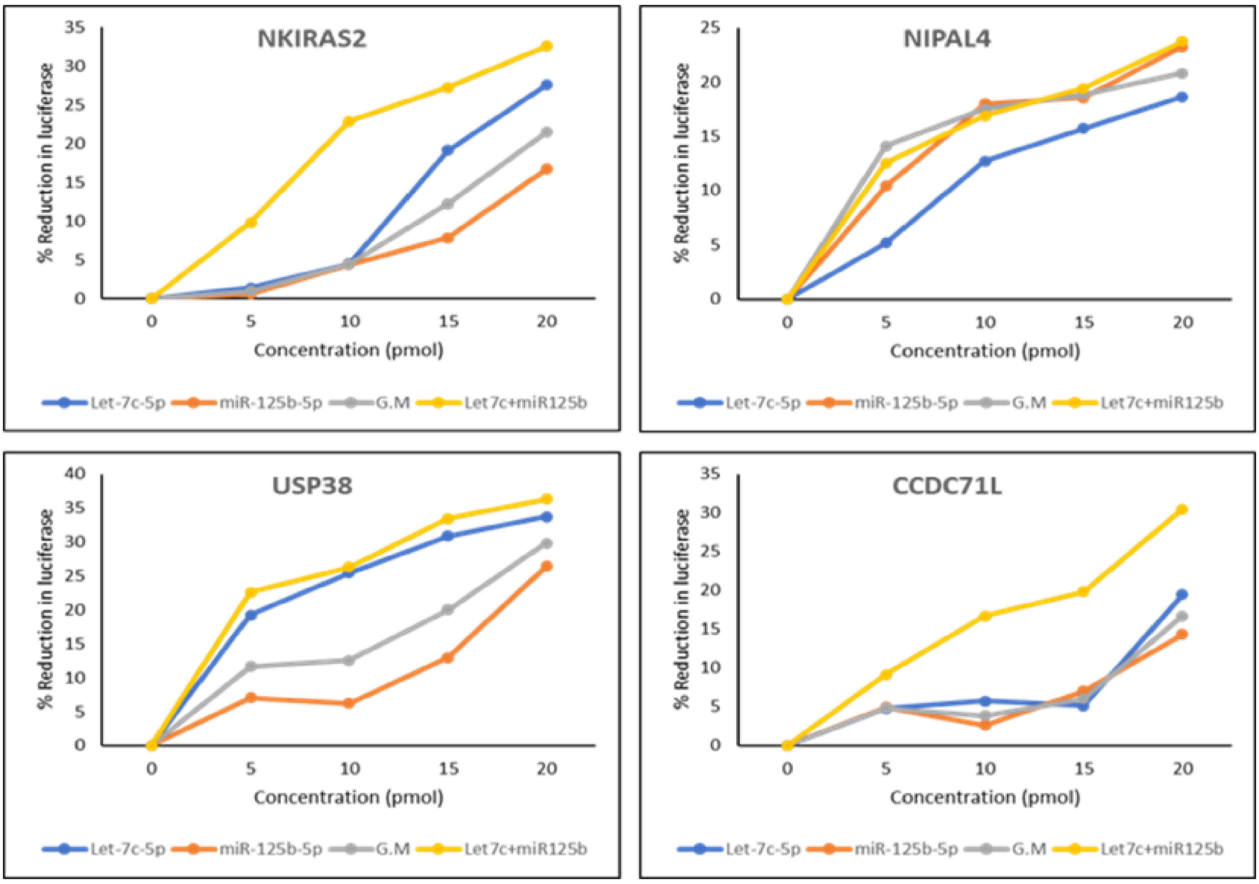
NKIRAS2
NIPAL4
USP38
CCDC71L
% Reduction in luciferase
Concentration (pmol)
0
5
10
15
20
25
30
35
40
Let-7c-5p
miR-125b-5p
G.M
Let7c+miR125b

### Figure S5

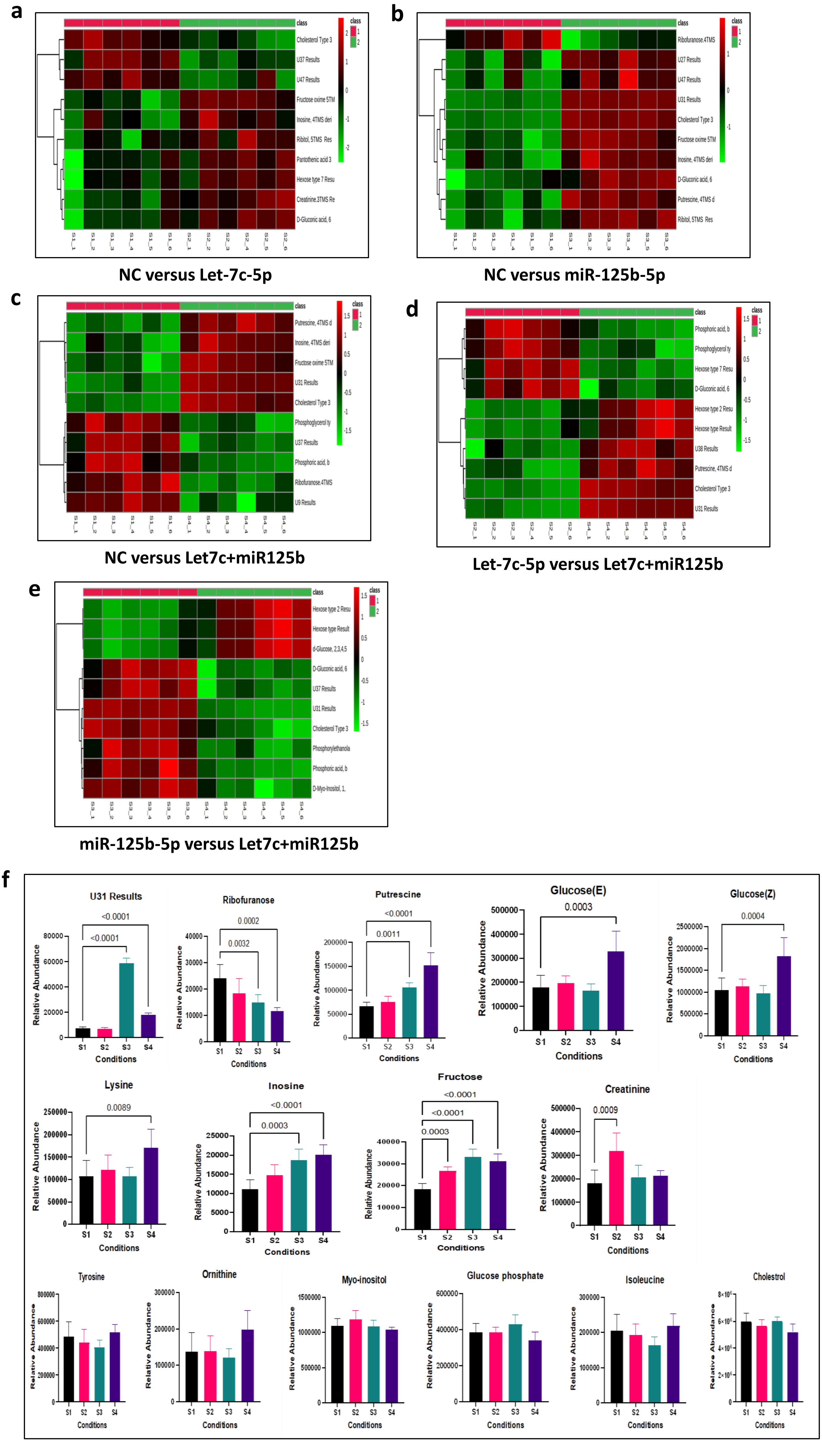

a
NC versus Let-7c-5p
b
NC versus miR-125b-5p
c
NC versus Let7c+miR125b
d
Let-7c-5p versus Let7c+miR125b
e
miR-125b-5p versus Let7c+miR125b
f
U31 Results
Ribofuranose
Putrescine
Glucose(E)
Glucose(Z)
Lysine
Inosine
Fructose
Creatinine
Tyrosine
Ornithine
Myo-inositol
Glucose phosphate
Isoleucine
Cholestrol
Relative Abundance
Conditions
S1
S2
S3
S4
<0.0001
0.0002
0.0032
0.0011
0.0003
0.0004
0.0089
0.0009
